# Diverse toxins exhibit a common binding mode to the Nicotinic Acetylcholine Receptors revealing a new molecular determinant

**DOI:** 10.1101/2024.06.04.597380

**Authors:** Hung N. Do, Jessica Z. Kubicek-Sutherland, S. Gnanakaran

## Abstract

Nicotinic acetylcholine receptors (nAChRs) are critical ligand-gated ion channels in the human nervous system. They are targets for various neurotoxins produced by algae, plants, and animals. While there have been many structures of nAChRs bound by neurotoxins published, the binding mechanism of toxins to the nAChRs remains uncleared. In this work, we have performed extensive Gaussian accelerated molecular dynamics simulations on several *Aplysia californica* (AC) nAChRs in complex with α-conotoxins, strychnine, and pinnatoxins, as well as human nAChRs in complex with α-bungarotoxin and α-conotoxin for a total of 60 μs of simulation time to determine the binding and dissociation pathways of the toxins to the nAChRs and the associated effects. We uncovered two common binding and dissociation pathways shared by toxins and nAChRs. In the primary binding pathway, the toxins diffused from the bulk solvent to first bind a region near the extracellular pore before moving downwards along the nAChRs to the nAChR orthosteric pocket. The second binding pathway involved a direct diffusion of the toxins from the bulk solvent into the nAChR orthosteric pocket. The dissociation pathways were the reverse of the observed binding pathways. We also found that the toxins enacted their toxicity upon binding by restricting the necessary movements required by the nAChRs to open their extracellular and intracellular pores for the ions to pass through. Notably, the electrostatically bipolar interactions between nAChR orthosteric pocket and toxins provides a molecular level explanation for the common binding mode shared by diverse toxins and serve as a key determinant for toxicity.

## Introduction

Nicotinic acetylcholine receptors (nAChRs) are critical ligand-gated ion channels in the human nervous system, which can be divided into two groups including muscle receptors and neuronal receptors, and targets for various neurotoxins produced by algae, plants, and animals^1,2^. nAChRs are pentameric (five subunits) proteins that can be composed of homomeric subunits (all of the same type such as α7, α8, and α9) and heteromeric subunits (with multiple subunits combined from the combinations of α2-α6 with β2-β4, α9 with α10, or α1 with β1, δ, and γ or ε), resulting in diverse functional properties^1,3^. A total of seventeen different nAChR subunits have been identified, including α1-α10, β1-β4, δ, γ and ε^1^. The subunits are structurally similar but diverse in sequences^3^, thus displaying varying binding interactions and affinities towards different toxins^4^. nAChRs can be referred to as a model of allosteric protein, with a full nAChR structure consisting of the extracellular ligand binding domain, a bundle of transmembrane helices, and a cytoplastic C-terminal latch that was shown to modulate channel opening^2,5^. The gating mechanism of nAChRs have been shown to consist of three primary states, including the resting, activated, and desensitized state^5^. In the resting state, an antagonist binds to the nAChR, resulting the closure of cationic channel of the receptor^5^. In the activated state, an agonist binds to the ligand binding pocket of the nAChR, opening the cationic channel and making the receptor highly permeable to cations^5^. The desensitized state could be considered the intermediate state between the resting and activated state, as the cationic channel of nAhCR closes to ion permeation despite the binding of an agonist^5^. Protein toxins produced by marine snails (α-conotoxins) and snakes (α-bungarotoxins) have been shown to display especially high affinity and selectivity for nAChRs^6^.

Conotoxins are a family of extremely toxic neurotoxins composed of cysteine-rich peptides produced by marine cone snails^7–9^. They are small 13-15 amino acid peptide toxins that act as competitive inhibitors of nAChRs binding to the acetylcholine ligand binding site located between subunits^10^. The natural conotoxin sequences are incredibly diverse^11^. It is estimated that less than two percent of existing conotoxins have been sequenced^12^. Over the last two decades, many X-ray crystal structures of α-conotoxins bound to the soluble acetylcholine binding protein (AChBP), a structural homolog of nAChR containing the ligand binding extracellular domain, have been published^10^. α-Bungarotoxin is a relatively large 74 amino acid peptide toxin found in the venom of certain snakes^13^. It has been shown to irreversibly bind to the human α7 nAChR subunit^14^, with a cryo-EM structure published^5^. Strychnine is a highly potent alkaloid (non-protein) neurotoxin produced by poisonous plants that has been shown to inhibit muscle and neuronal nAChRs^15^. A X-ray crystal structure of strychnine in complex with AChBP has been published^16^. Pinnatoxin is a macrocyclic imine (non-protein) marine phycotoxin produced by algae with high affinity for α7 and α1_2_βγδ nAChRs^17^. X-ray crystal structures of pinnatoxin A and pinnatoxin G in complex with AChBP have been published^17^.

Molecular dynamics (MD) is a compelling computational technique for simulating biomolecular dynamics on an atomistic level^18^ and can lead to the understanding of binding modes. Gaussian accelerated molecular dynamics (GaMD) is an unconstrained enhanced sampling technique that facilitates dynamic transitions by applying a harmonic boost potential to smooth the biomolecular potential energy surface^19^. Since the boost potential usually exhibits a near Gaussian distribution, GaMD can maintain the overall shape of the original free energy surface, and cumulant expansion to the second order (“Gaussian approximation”) can be applied to achieve proper energy reweighting^19^. GaMD has been demonstrated to properly sample the biological processes of ligand binding^20^ and protein conformational changes upon ligand binding^21^.

In this work, we want to examine whether a diverse set of toxins (α-conotoxins, strychnine, and pinnatoxins) (**Figure 1**) share a common binding mode regardless of the origin (human or sea organism) or the different constructs (soluble surrogate and the fully membrane-bound) of nAChR. We performed GaMD simulations on a number of *Aplysia californica* (AC) nAChRs in complex with α-conotoxins, strychnine, and pinnatoxins, as well as human membrane-bound nAChRs in complex with α-bungarotoxin and α-conotoxin (**Figures 2** and **S1**) to determine the binding and dissociation pathways of the toxins to the nAChRs as well as the associated effects. It is our understanding that AC nAChRs were chosen to model human nAChRs due to their high similarity to the α7 nAChR^22^. We deduce a molecular explanation for the common binding mode shared across multiple ligands in the nAChRs. Our work provides important mechanistic insights into the binding of toxins in the nAChRs and identifies key binding descriptors that could be useful for machine learning predictive models.

**Figure 1.**
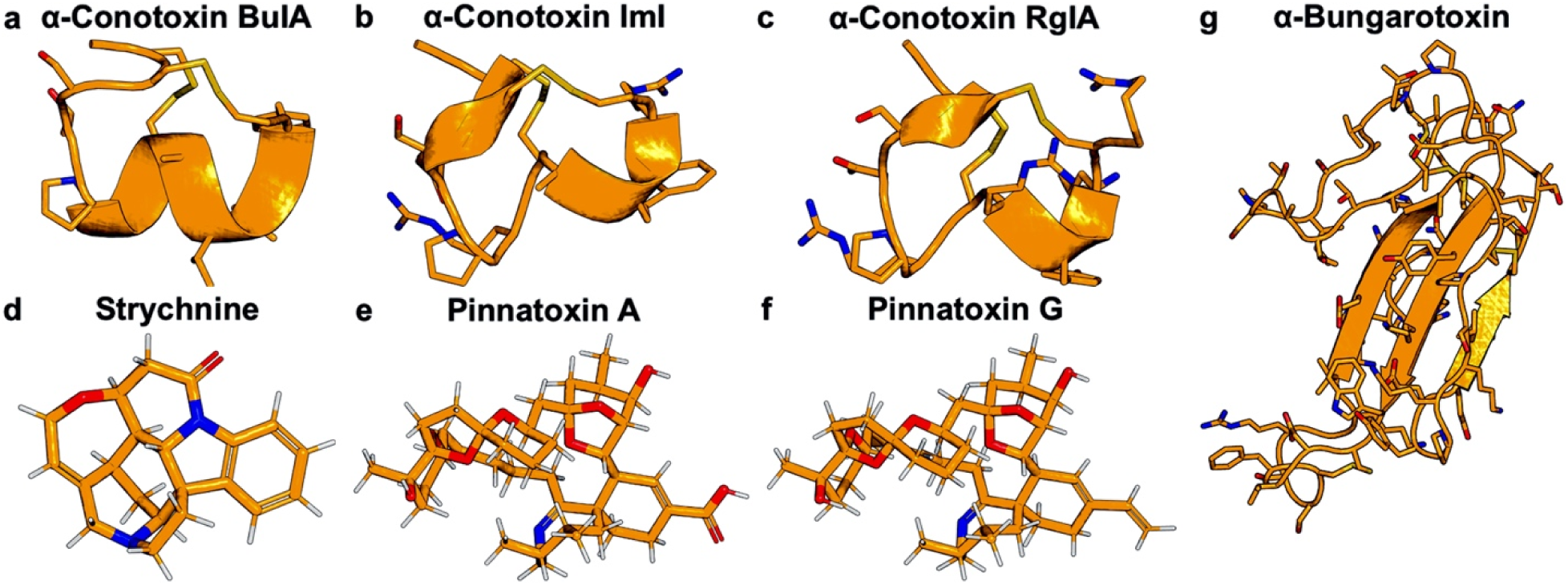
Structures of toxins explored in this study, including **(a)** α-conotoxin BuIA, **(b)** α-conotoxin ImI, **(c)** α-conotoxin RgIA, **(d)** strychnine, **(e)** pinnatoxin A, **(f)** pinnatoxin G, and **(g)** α-bungarotoxin. The heteroatoms shown as sticks are colored as following: nitrogen (blue), oxygen (red), hydrogen (white), and sulfur (dark yellow). The PDB IDs of the structures are included in **Table S1**.

**Figure 2.**
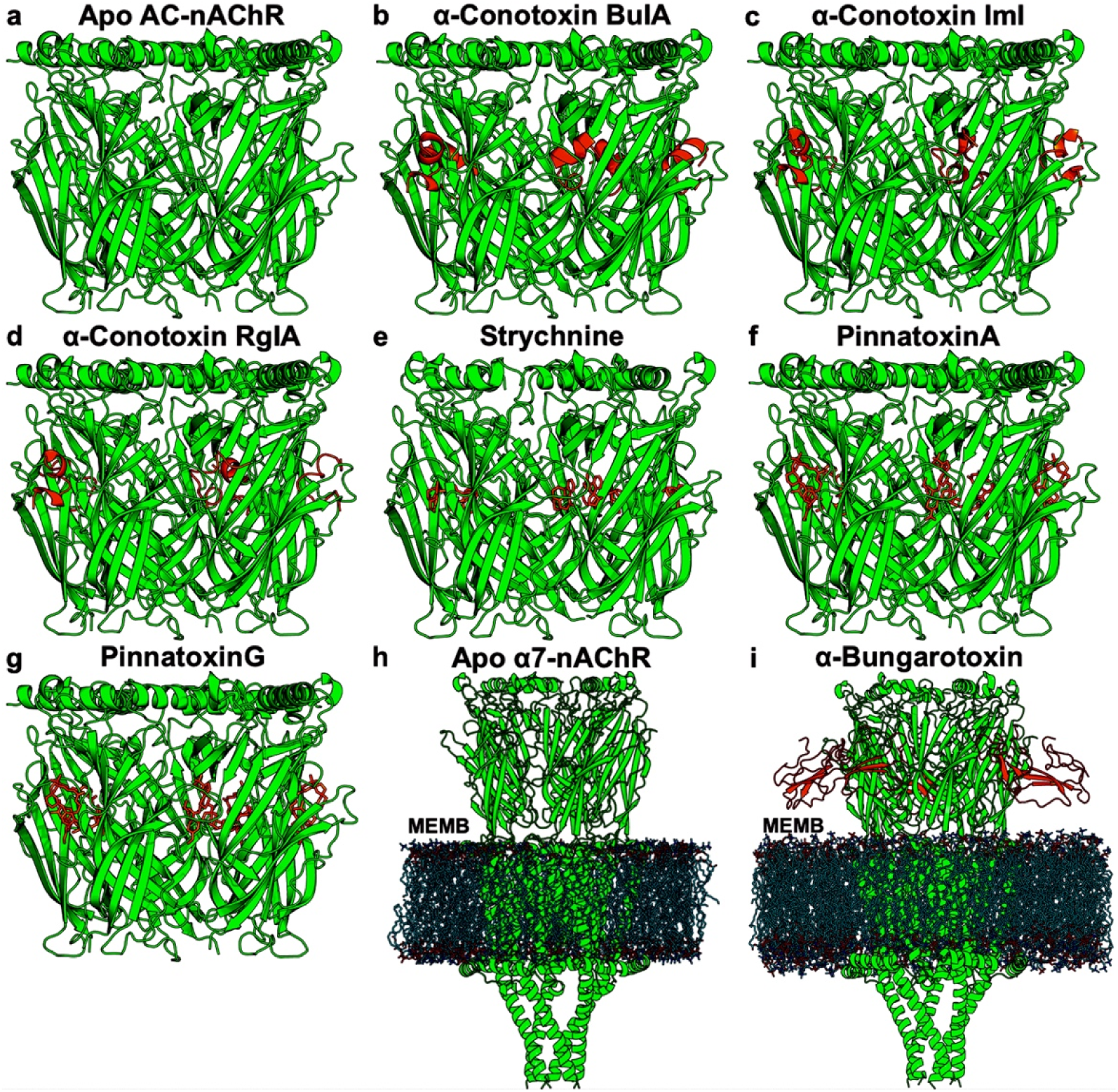
Simulation systems of nicotinic acetylcholine receptors (nAChRs) bound by toxins. **(a)** Apo Aplysia californica (AC) nAChR. **(b)** AC nAChR bound by α-conotoxin BuIA. **(c)** AC nAChR bound by α-conotoxin ImI. **(d)** AC nAChR bound by α-conotoxin RgIA. **(e)** AC nAChR bound by strychnine. **(f)** AC nAChR bound by pinnatoxin A. **(g)** AC nAChR bound by pinnatoxin G. **(h)** Apo human α7 nAChR. **(i)** Human α7 nAChR bound by α-bungarotoxin. The nAChRs are colored green, while the toxins are colored orange. The POPC membrane lipid bilayers were shown in **(h-i)**. Additional simulation systems of toxin binding to the nAChRs were included in **Figure S1**.

## Methods

### Simulation system setup

We started from the X-ray crystal structures of *Aplysia californica* (AC) nAChR in apo state (PDB: 2BYN)^22^ and in complex with α-conotoxin BuIA (PDB: 4EZ1)^23^, α-conotoxin ImI (PDB: 2BYP)^22^, α-conotoxin RgIA (PDB: 7EGR)^24^, strychnine (PDB: 2XYS)^16^, pinnatoxin A (PDB: 4XHE)^17^, and pinnatoxin G (PDB: 4XK9)^17^, and the cryo-EM structures of human α7 nAChR in apo state (PDB: 7EKI)^25^ and in complex with α-bungarotoxin (PDB: 7KOO)^26^ to prepare the simulation systems (**Figures 1-2** and **Table S1**). The missing residues within the structures were restored using the SWISS-MODEL homology modeling webserver^27^, including residues P18-M19 in the 2BYN^22^, F14-M19 in the 4EZ1^23^, and S17-M19 in the 2BYP^22^ PDB structure. The starting structures of the AC nAChR, human α7-nAChR, and human α3β4 nAChR with free α-conotoxin RgIA were prepared by removing the orthosteric ligands (if present) from the nAChR orthosteric pockets in the 2BYN^22^, 7EKI^25^, and 6PV7^28^ PDB structures, respectively, and placing five RgIA molecules ∼20 Å away from the receptors extracellularly (**Figure S1** and **Table S1**). The CHARMM-GUI webserver^29–31^ was used to prepare the simulation systems. All heteroatom molecules, except the toxins, were removed from the starting structures. The protein chain termini were capped with neutral patches (acetyl and methylamide), except the C-terminal of the α-conotoxin BuIA, where amide was used to be consistent with the 4EZ1^23^ PDB structure. The starting structures of AC nAChR simulation systems were solvated in 0.15 M NaCl solution boxes that extended ∼10 Å from the solute surfaces. The structures of human nAChR simulation systems were embedded in POPC membrane lipid bilayers and then solvated in 0.15 M NaCl solutions. The AMBER^32^ force field parameter sets were used for the GaMD simulations, including the ff14SB^33^ for proteins, GAFF2^34^ for ligands using the AM1-BCC^35^ charging method, LIPID17 for lipids, and TIP3P^36^ for water.

### Simulation protocols

All-atom dual-boost GaMD simulations^19^ were performed on the nAChR simulation systems to examine the binding and dissociation of the toxins to the receptors. Periodic boundary conditions were applied to the simulation systems, and bonds containing hydrogen atoms were restrained with the SHAKE^37^ algorithm. A timestep of 2 fs was used. The temperature was kept constant at 310 K using the Langevin thermostat^38,39^ with a friction coefficient of 1.0 ps^-1^. The electrostatic interactions were calculated using the particle mesh Ewald (PME) summation^40^ with a cutoff of 9.0 Å for long-range interactions. The pressure was kept constant at 1.0 bar using the Berendsen barostat^41^ with isotropic coupling for the soluble AC nAChR simulation systems and with semi-isotropic coupling with enabled surface tension in the X-Y plane for the membrane-bound human nAChR simulation systems, respectively. The Berendsen coupling constant was set to 0.5 ps. The GPU version of the *pmemd* module (*pmemd.cuda*) in AMBER 22^42^ was used to carry out the MD simulations of nAChRs. The simulation systems were energetically minimized for 5000 steps using the steepest-descent algorithm and equilibrated with the constant number, volume, and temperature (NVT) ensemble. The membrane-bound human nAChR systems were further equilibrated with the constant number, pressure, and temperature (NPT) ensemble. The conventional MD (cMD) simulations were then performed for 10 ns using the NPT ensemble. GaMD simulations of the soluble AC nAChR systems involved an initial short cMD simulation of 4-10 ns to calculate GaMD acceleration parameters, followed by the GaMD equilibration stage of 16-40 ns to equilibrate and update the acceleration parameters. On the other hand, GaMD simulations of the membrane-bound human nAChR systems consisted of an initial short cMD simulation of 8-10 ns and GaMD equilibration simulation of 32-40 ns. Five independent 1000 ns GaMD production simulations with randomized initial atomic velocities and acceleration parameters kept constant were performed for each nAChR system. The dual-boost scheme was used, with one boost potential applied to the dihedral energetic term and another to the total potential energetic term. The reference energy *E* was set to lower bound (*E* = *V_max_*) for both dihedral and total potential energy. The upper limits of the boost potential standard deviations, σ_0_, were set to 6.0 kcal/mol for both the dihedral (σ_0D_) and total (σ_0P_) potential energetic terms. The GaMD simulations are summarized in **Table S1**.

### Simulation analysis

The CPPTRAJ^43^ tool was used to analyze the simulation trajectories. The PyReweighting^44^ toolkit was used to calculate the potential mean force (PMF) free energy profiles using the center-of-mass (COM) distance between strychnine as well as the Cα atoms of α-conotoxin RgIA and the nAChR orthosteric pockets (including residues Q186 – Y195 in all five subunits of the AC nAChR, residues R185 – Y194 in all five subunits of the human α7-nAChR, and residues A:I188 – A:Y197, B:T190 – B:D200, C:T190 – C:D200, D:I188 – D:Y197, and E:T190 – E:D200 in the human α3β4-nAChR) and the nAChR pore diameters (Cα-atom distances between residues A:Q100 and D:Q100 in the AC nAChR, A:A101 and D:A101 in the human α7-nAChR, and A:D103 and C:T103 in the human α3β4-nAChR). The numbering scheme of the AC nAChR is based on the 2BYP^22^ PDB, of the human α7 nAChR is based on the 7KOO^26^ PDB, and of the human α3β4 nAChR is based on the 6PV7^28^ PDB structure, with the chain ID listed before the colon and residue numbers included after the colons. A bin size of 1.0-2.0 Å and cutoff of 100 frames in one bin were used for reweighting of the GaMD simulations of strychnine dissociation and RgIA binding in the nAChRs. Furthermore, we considered nAChR residues within 4 Å distance of the toxins to be interacting residues with the ligands. The hierarchical agglomerative clustering algorithm was used to obtain representative conformations of the low-energy conformational states observed in the binding and dissociation of toxins in the nAChRs. Lastly, we also calculated the root-mean-square fluctuations (RMSFs) and performed principal component analysis (PCA) to determine the changes in nAChR flexibility upon toxin binding and primary motions in the simulation systems, respectively, to examine the effects of toxin binding to the nAChRs. In **Figure 5**, only the motions with ≥6Å distances with RMSDs of ∼3Å corresponding to the first principal components were shown for all simulation systems.

## Results

All-atom dual-boost GaMD simulations were performed on the apo AC nAChR, α-conotoxin-BuIA-bound AC nAChR, α-conotoxin-ImI-bound AC nAChR, α-conotoxin-RgIA-bound AC nAChR, strychnine-bound AC nAChR, pinnatoxin-A-bound nAChR, pinnatoxin-G-bound nAChR, AC nAChR with free α-conotoxin RgIA, apo human α7 nAChR, α-bungarotoxin-bound human α7 nAChR, human α7 nAChR with free α-conotoxin RgIA, and human α3β4 nAChR with free α-conotoxin RgIA (**Figures 1-2** and **S1**). The seven toxins explored in this study illustrate various physical and structural properties – three peptides (α-conotoxins BuIA, ImI, and RgIA), one protein (α-bungarotoxin), one small chemical compound (strychnine), and two large chemical compounds (pinnatoxin A and G) – as well as originate from different sources – three from sea snails (α-conotoxins BuIA, ImI, and RgIA), one from snakes (α-bungarotoxin), one from poisonous plants (strychnine), and two from algae (pinnatoxin A and G) (**Figure 1**). In addition, the constructs of the nAChRs were also diverse: the AC nAChRs only included the soluble extracellular ligand binding domain, whereas the human α7 and α3β4 consisted of the soluble extracellular ligand binding domains, the transmembrane helical regions, as well as the intracellular domains (**Figures 2** and **S1**).

The boost potentials applied in the GaMD simulations of the nAChRs were recorded to be similar across the systems, with the average values of 11.8 – 14.6 kcal/mol and standard deviations of 3.8 – 4.6 kcal/mol (**Table S1**). Notably, GaMD simulations captured dissociation of strychnine from the AC nAChR (**Figures 2** and **S3**) and multiple binding and dissociation events of α-conotoxin RgIA to the AC nAChR (**Figures 3** and **S5**), the human α7 nAChR (**Figures S6** and **S8**), and the human α3β4 nAChR (**Figure S9**). Overall, the dissociation pathway observed for strychnine in the AC nAChR was the reverse of the primary binding pathway observed for α-conotoxin RgIA in the AC nAChR, human α7 nAChR, and human α3β4 nAChR. Furthermore, it was observed that different toxins of different sizes and chemical properties bound to the nAChRs of different organisms and subunits using the identical pathways.

**Figure 3.**
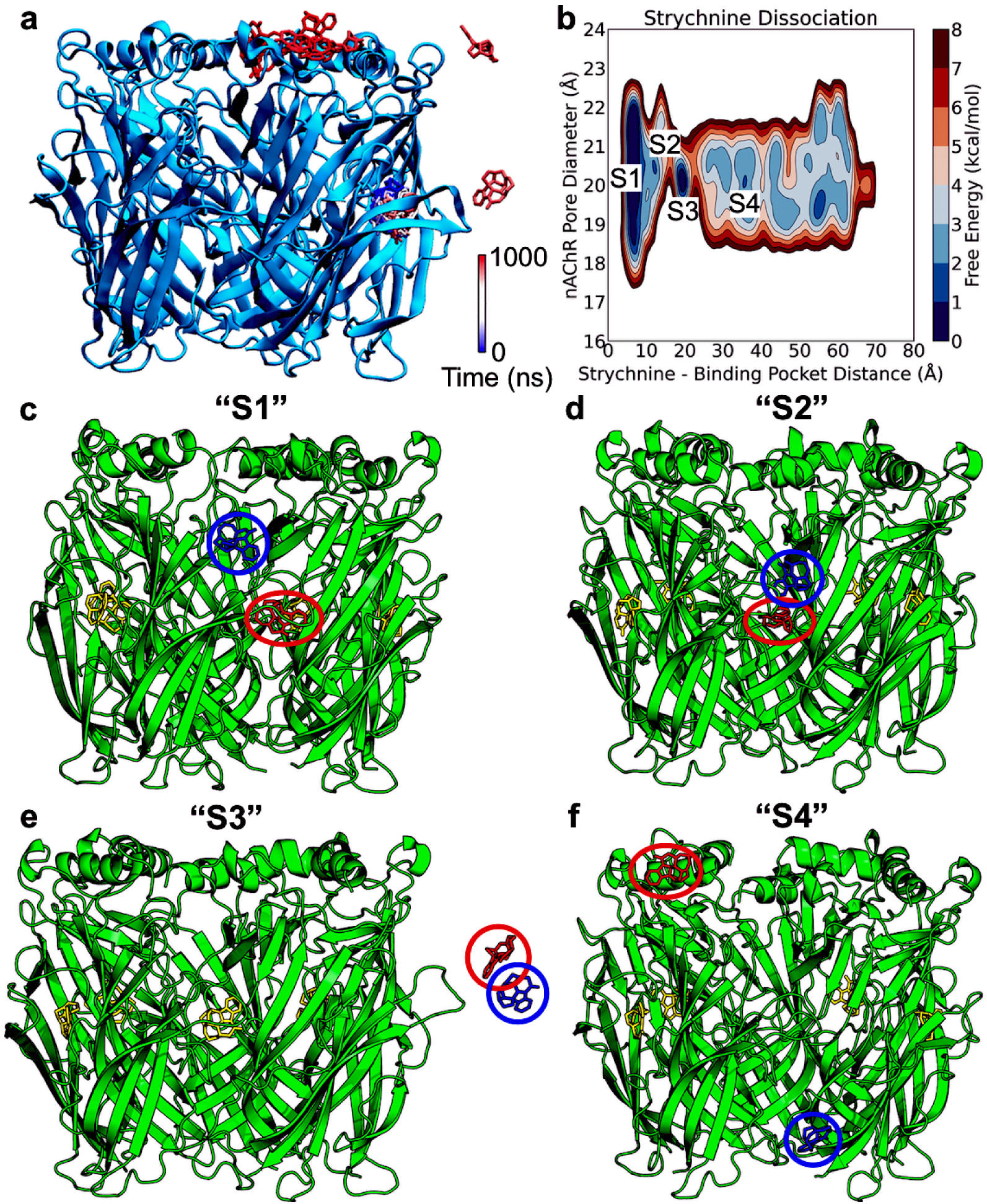
Dissociation of strychnine from the Aplysia californica nicotinic acetylcholine receptor. **(a)** Trace of the strychnine molecule dissociated from the nAChR observed in Sim4. A color scale of blue (0 ns) – white – red (1000 ns) is used to show the location of strychnine during Sim4. **(b)** 2D potential mean force (PMF) free energy profile of the strychnine dissociation from the nAChR calculated from the COM distance between strychnine and Cα atoms of the orthosteric binding pocket of nAChR (consisting of residues E:Q186 to E:Y195) and the nAChR pore diameter (Cα-distance between residues A:Q100 and D:Q100). Selected low-energy conformational states are labeled “S1”–“S4”. **(c)** The “S1” state, where the distance between strychnine and nAChR orthosteric pocket is ∼6.9 Å and the nAChR pore diameter is ∼20.8 Å. **(d)** The “S2” state, where the distance between strychnine and nAChR orthosteric pocket is ∼13.2 Å and the nAChR pore diameter is ∼20.9 Å. **(e)** The “S3” state, where the distance between strychnine and nAChR orthosteric pocket is ∼20.4 Å and the nAChR pore diameter is ∼19.8 Å. **(f)** The “S4” state, where the distance between strychnine and nAChR orthosteric pocket is ∼35.2 Å and the nAChR pore diameter is ∼20.4 Å. The strychnine-1 is colored and circled in red, strychnine-2 is colored and circled in blue, while the other strychnine molecules are colored yellow.

### Dissociation of the small chemical toxin strychnine from the *Aplysia californica* nAChR

Initially, six molecules of strychnine bound to the AC nAChR, with four orthosteric pockets of nAChR accommodating one strychnine molecule each and one orthosteric pocket containing two strychnine molecules (**Figure 2e**). For the purpose of this study, we will refer to the two strychnine molecules that bound the same nAChR orthosteric pocket as strychnine-1 and strychnine-2, with strychnine-1 being the inner strychnine molecule that bound the orthosteric pocket more tightly and strychnine-2 being the outer strychnine molecule (**Figure 2e**).

The dissociation of strychnine-1 from nAChR orthosteric pocket observed only once in Sim4 was shown in **Figure 3a** and facilitated by strychnine-1 interactions with strychnine-2 (**Figures 3, S2,** and **S3**). A two-dimensional (2D) potential mean force (PMF) free energy profile was calculated from the COM distance between strychnine-1 and the nAChR orthosteric pocket (including residues E:Q186 – E:Y195) and the nAChR pore diameter (Cα-atom distance between residues A:Q100 and D:Q100) in Sim4 to characterize strychnine-1 dissociation from the AC nAChR (**Figure 3b**). We selected four low-energy conformational states from the 2D PMF free energy profile to describe strychnine-1 dissociation from the nAChR, namely states “S1” – “S4”. In the “S1” low-energy conformational state, the distance between strychnine-1 and nAChR orthosteric pocket was ∼6.9 Å (**Figure 3c**). Strychnine-1 was located inside the nAChR orthosteric pocket, interacting with residues A:Y55, E:Y93, E:S146-E:W147, and residues E:Y188, E:C190-E:C191, and E:Y195 of the conserved Y-CC-EPY motif in the nAChR orthosteric pocket. Meanwhile, strychnine-2 was located upwards nearby, interacting with residues E:K25, E:P28, and E:G151-E:F152. In state “S2”, the distance between strychnine-1 and nAChR orthosteric pocket was ∼13.2 Å (**Figure 3d**). In this state, strychnine-2 was observed to go back inside the nAChR orthosteric pocket and interact with strychnine-1 to drag strychnine-1 out of the orthosteric pocket (**Figure 3d**), which resulted in the strychnine-1 dissociation observed in the “S3” low-energy conformational state (**Figure 3e**). In the “S3” state, strychnine-1 was observed to be floating in the bulk solvent while interacting with strychnine-2. Finally, the two toxin molecules broke apart in state “S4”, where the distance between strychnine-1 and the nAChR orthosteric pocket was ∼35.2 Å (**Figure 3f**). Strychnine-1 then moved up to the extracellular pore of the nAChR, interacting with residues R843, M893, while strychnine-2 bound to a pocket outside the intracellular pore in another subunit of the nAChR, interacting with residues A:L165, A:Y168-A:S172, A:Y174-A:I176, and A:R207. Overall, strychnine-1 was able to dissociate from the nAChR mainly because of its interactions with the flexible strychnine-2, which dragged the toxin molecule outwards. Strychnine-1 then moved and bound to the extracellular pore of the nAChR (**Figure 3a**).

Unlike the other five strychnine molecules that bound tightly to the AC nAChR (**Figure S2**), strychnine-2 binding to the AC nAChR was unstable. In particular, the molecule was observed to dissociate from the nAChR orthosteric pocket in all five simulations (**Figure S3a**). The dissociation pathway of strychnine-2 (**Figure S3b**) was observed to be different from that of strychnine-1 (**Figure 3a**). From its initial position in the nAChR orthosteric pocket, interacting with residues A:T36, A:Q57, A:M116, A:I118, A:D164, and residues E:C190-E:C191 of the conserved Y-CC-EPY motif in the nAChR orthosteric pocket, strychnine-2 fluctuated and gradually diffused outside of the nAChR orthosteric pocket into the bulk solvent (**Figure S3b**). The phenomenon that two ligand molecules binding to the same orthosteric pocket showed different binding and dissociation pathways has been observed in a previous study^20^.

### Association of the α-conotoxin RgIA to the *Aplysia californica* nAChR

We captured two different binding pathways in a total of seven binding events across all five simulation replicas (**Figure S4**) of the α-conotoxin RgIA to the AC nAChR orthosteric pocket in the simulation system of AC nAChR with free α-conotoxin RgIA (**Figure 1h**). The primary binding pathway was mostly the reverse of the dissociation pathway of strychnine-1 from the nAChR (**Figure 3a**) and described in **Figure 4**. The secondary binding pathway was a direct binding of the α-conotoxin RgIA from the bulk solvent into the nAChR orthosteric pocket (**Figure S5**), which was the reverse of the dissociation pathway of strychnine-2 from the nAChR (**Figure S3b**).

**Figure 4.**
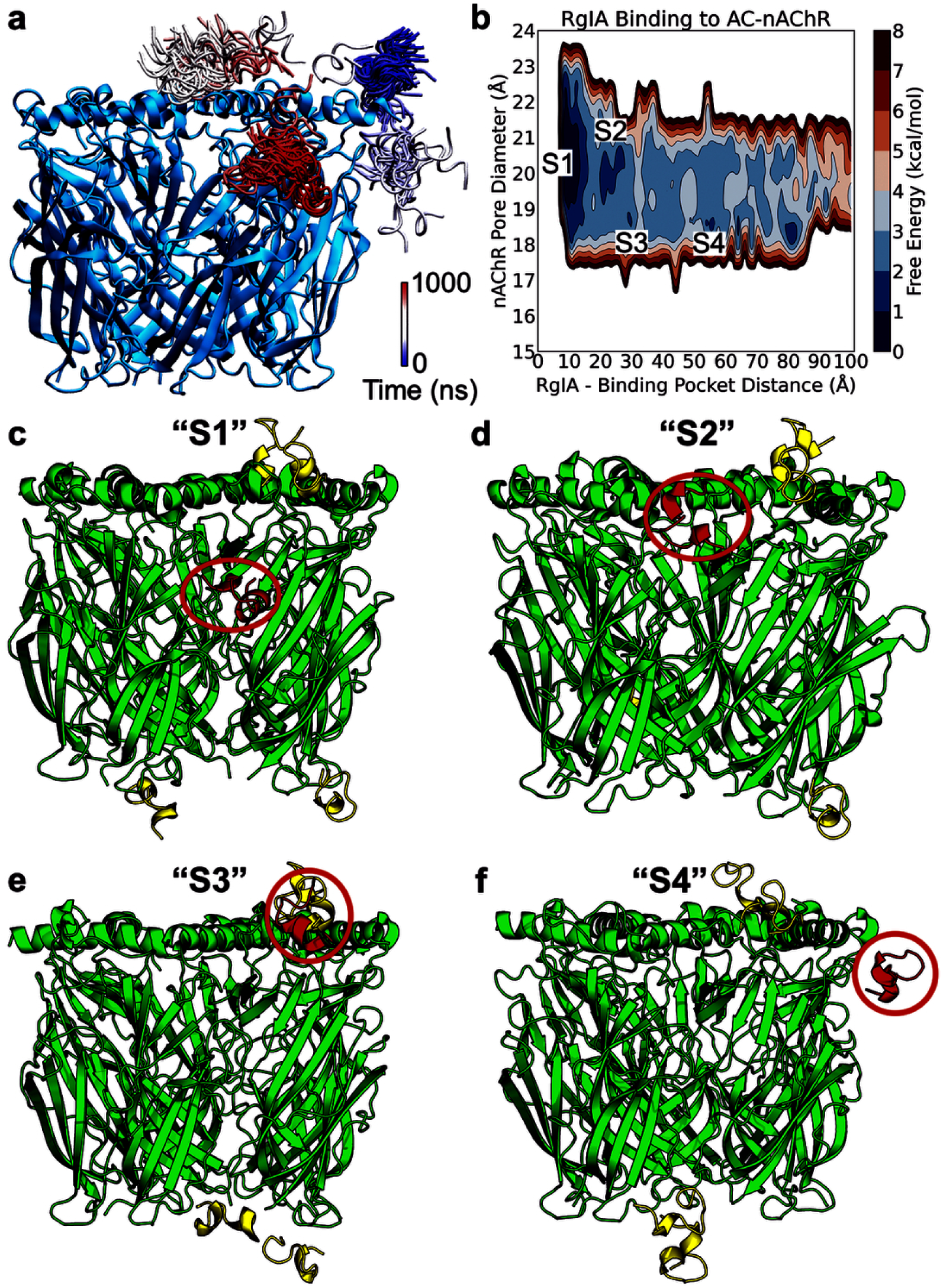
Binding of α-conotoxin RgIA to the Aplysia californica nicotinic acetylcholine receptor. **(a)** Representative trace of α-conotoxin RgIA binding to the AC nAChR observed in Sim2. A color scale of blue (0 ns) – white – red (1000 ns) is used to show the location of α-conotoxin RgIA during Sim2. **(b)** 2D potential mean force (PMF) free energy profile of the α-conotoxin RgIA binding to the AC nAChR calculated from the COM distance between Cα atoms of RgIA and the orthosteric binding pockets of AC nAChR (consisting of residues Q186 – Y195 in the subunits) and the nAChR pore diameter (Cα-distance between residues A:Q100 and D:Q100). Selected low-energy conformational states are labeled “S1”–“S4”. **(c)** The “S1” state, where the distance between RgIA and nAChR orthosteric pocket is ∼9.1 Å and the nAChR pore diameter is ∼20.8 Å. **(d)** The “S2” state, where the distance between RgIA and nAChR orthosteric pocket is ∼19.8 Å and the nAChR pore diameter is ∼20.7 Å. **(e)** The “S3” state, where the distance between RgIA and nAChR orthosteric pocket is ∼30.5 Å and the nAChR pore diameter is ∼19.0 Å. **(f)** The “S4” state, where the distance between RgIA and nAChR orthosteric pocket is ∼53.7 Å and the nAChR pore diameter is ∼19.1 Å. The α-conotoxin RgIA of interest is colored and circled in red, while the other α-conotoxin RgIA molecules are colored yellow.

A 2D PMF free energy profile was calculated from the COM distances between Cα atoms of the α-conotoxin RgIA and the AC nAChR orthosteric pockets (including residues Q186-Y195 in the subunits) and the nAChR pore diameters (Cα-atom distance between residues A:Q100 and D:Q100) in the simulations in which RgIA was observed to bind the AC nAChR (**Figure S4**) to characterize the primary binding pathway of α-conotoxin RgIA to the nAChR orthosteric pocket (**Figure 4b**). Four different low-energy conformational states, namely “S1”-“S4”, were selected to describe this first pathway. In the “S1” state, the distance between RgIA and nAChR orthosteric pocket was ∼9.1 Å. The α-conotoxin bound to the nAChR orthosteric pocket in this state, interacting with residues B:D77, B:R79, C:S150, C:F152-C:E153, and residues C:Y188-C:E193 and C:Y195 of the conserved Y-CC-EPY motif in the nAChR orthosteric pocket (**Figure 4c**). In the “S2” low-energy conformational state, the distance between RgIA and nAChR orthosteric pocket was ∼19.8 Å. In this state, the α-conotoxin of interest was located at the interface between two subunits near the extracellular pore directly above the orthosteric pocket of AC nAChR, interacting with nAChR residues B:D(−2), B:N70- B:E71, B:G73-B:N74, and C:K25-C:D26 (**Figure 4d**). In the “S3” and “S4” low-energy state, the α-conotoxin RgIA of interest was located near the extracellular pore of a nearby subunit (**Figure 4e-4f**). In the “S3” low-energy state, the α-conotoxin was interacting with nAChR residues B:D68 and B:N70 (**Figure 4e**), while in the “S4” state, the α-conotoxin was interacting with nAChR residues A:P69-A:N70, A:N74, A:T76- A:D77, A:T110, B:K25-B:D26, and B:F152 (**Figure 4f**). Overall, the primary binding pathway of α-conotoxin RgIA to the nAChR involved its movement from the nAChR extracellular pore to the nAChR orthosteric pocket (**Figure 4a**), which was the reverse of the strychnine-1 dissociation pathway (**Figure 3a**). Meanwhile, in the secondary binding pathway observed in Sim1, the α-conotoxin RgIA diffused from the bulk solvent directly into the nAChR orthosteric pocket (**Figure S5**), the reverse of the dissociation pathway of strychnine-2 (**Figure S3b**).

### Common binding pathways of toxins to the nAChRs shared across toxins, organisms, and subunits

In the GaMD production simulations of human α7 and α3β4 with free α-conotoxin RgIA (**Figure S1b-S1c**), we observed a total of eight and four distinct binding events, respectively, across four out of five simulation replicas for each simulation system (**Figures S6-S7**). Overall, the binding pathways of the α-conotoxin RgIA to human α7 and α3β4 nAChRs were highly similar to the other (**Figures S8-S10**) and also almost identical to the binding pathway of the α-conotoxin RgIA to the AC nAChR (**Figures 4** and **S5**). Notably, binding of the α-conotoxin RgIA to human α7 and α3β4 nAChRs only involved the soluble extracellular ligand binding domains and did not pertain to either the transmembrane helical regions or the intracellular domains (**Figures S8-S10**). Since we captured fewer binding events of α-conotoxin RgIA to the human α3β4 nAChR compared to the human α7 nAChR, the toxin appeared to be selective towards the human α7 nAChR^45^.

The α-conotoxin RgIA was first observed to bind the human α7 nAChR in the 50 ns GaMD equilibration simulation (**Figure S8a**). This binding pathway closely resembled the primary binding pathway of the toxin to the AC nAChR (**Figure 4a**), in which RgIA moved from the bulk solvent to first bind the nAChR extracellular pore, then gradually diffused outwards, and moved to the nAChR orthosteric pocket (**Figure S8a**). Furthermore, another RgIA peptide was also observed to bind to the human α7 nAChR in Sim2 in a similar manner (**Figure S8b**). Meanwhile, the α-conotoxin RgIA that bound to the nAChR during the GaMD equilibration was observed to dissociate from the receptor in Sim2 and Sim5 in two different pathways (**Figure S8b**). In Sim2, the toxin dissociated from the human α7 nAChR in a pathway that was the reverse of the primary binding pathway of α-conotoxin RgIA to the nAChR, whereas in Sim5, the toxin diffused directly from the nAChR orthosteric pocket into the bulk solvent, the reverse of the secondary binding pathway of α-conotoxin RgIA to the nAChR (**Figure S8b**).

A slightly different binding pathway of the α-conotoxin RgIA to the human α7 nAChR was observed in Sim1 and described in **Figure S9**. This pathway could be referred to as a continuation of the primary binding pathway of α-conotoxin RgIA to the nAChR (**Figure S9a**). A 2D PMF free energy profile was calculated from the COM distance between Cα atoms of the α-conotoxin RgIA and the human α7 nAChR orthosteric pocket (consisting of residues R185 – Y194 in the subunits) and the nAChR pore diameter (Cα-atom distance between residues A:A101 and D:A101) in the simulations in which RgIA was observed to bind the human α7 nAChR (**Figure S6**) to characterize this binding pathway of the α-conotoxin RgIA to the human α7 nAChR orthosteric pocket (**Figure S9b**). We selected four low-energy conformational states, namely “S1”-“S4”, to characterize the binding of α-conotoxin RgIA to the human α7 nAChR observed in Sim1. In the “S1” low-energy state, the distance between RgIA and the nAChR orthosteric pocket was ∼14.1 Å. The RgIA of interest bound to the nAChR orthosteric pocket and interacted with nAChR residues A:Y31, A:Q56, A:S58, A:T60, A:Q65, A:N110, A:S113, A:H114, A:Q160, and residues E:Y187-E:E188 and E:C190-E:K191 of the conserved Y-CC-EPY motif (**Figure S9c**). In state “S2”, the distance between RgIA and the nAChR orthosteric pocket was ∼18.3 Å. The RgIA of interest interacted with residues A:Y31, A:Q56-A:M57, A:H114, A:Q116, A:Q158-A:E161, and E:E188 from the nAChR (**Figure S9d**). In the “S3” low-energy state, the distance between RgIA and the nAChR orthosteric pocket was ∼24.7 Å. The toxin was in contact with nAChR residues A:Y31, A:D156, and A:Q158-A:E161 (**Figure S9e**). The “S4” low-energy conformational state recorded a RgIA-nAChR orthosteric pocket distance of ∼39.4 Å. The RgIA toxin of interest interacted with residues A:S25, A:W153, A:D156, and A:E184 from the nAChR (**Figure S9f**). Overall, in this binding pathway, the α-conotoxin RgIA moved between two nAChR orthosteric pockets in two different subunits. The toxin first dissociated from the orthosteric pocket in the first nAChR subunit and then moved mostly horizontally to the orthosteric pocket in the second nAChR subunit (**Figure S9a**).

Finally, the primary binding pathway of the α-conotoxin RgIA to human α3β4 nAChR (**Figure S10a**) was highly similar to the primary binding of the α-conotoxin RgIA to the AC nAChR (**Figure 4a**). In particular, the toxin was first in contact with the α3β4 nAChR near the extracellular pore and gradually moved downwards along the nAChR towards the orthosteric pocket. A 2D PMF free energy profile was calculated from the COM distance between Cα atoms of the α-conotoxin RgIA and the human α3β4 nAChR orthosteric pocket (consisting of residues A:I188 – A:Y197, B:T190 – B:D200, C:T190 – C:D200, D:I188 – D:Y197, and E:T190 – E:D200) and the nAChR pore diameter (Cα-atom distance between residues A:D103 and C:T103) in the simulations in which RgIA was observed to bind the human α3β4 nAChR (**Figure S7**) to characterize this binding pathway of the α-conotoxin RgIA to the human α3β4 nAChR orthosteric pocket (**Figure S10b**). Three low-energy conformational states, including “S1”-“S3”, were selected to characterize the binding of the α-conotoxin RgIA to the human α3β4 nAChR. In the “S1” low-energy state, the distance between RgIA and the nAChR orthosteric pocket was ∼9.3 Å. In this state, the RgIA of interest bound to the nAChR orthosteric pocket and interacted with nAChR residues A:E4, A:E76, E:R24, E:S30, E:L32, E:H157-E:E159, E:V191-E:Q194, and E:S197 (**Figure S10c**). In the “S2” low-energy conformational state, the distance between RgIA and the nAChR orthosteric pocket was ∼15.0 Å. The RgIA toxin interacted with residues E:R24, E:S30, and E:T158-E:E159 from the α3β4 nAChR (**Figure S10d**). Finally, in state “S3”, the RgIA of interest was located at ∼20.1 Å distance from the nAChR orthosteric pocket and interacted with nAChR residues E:R24, E:S29-E:L32, and E:T158-E:E159 (**Figure S10e**).

### Effects of toxin binding to nAChRs

For other simulation systems of the nAChR, including the AC nAChR bound by α-conotoxin BuIA (**Figure 2b**), α-conotoxin ImI (**Figure 2c**), α-conotoxin RgIA (**Figure 2d**), pinnatoxin A (**Figure 2f**), and pinnatoxin G (**Figure 2g**), as well as the human α7 nAChR bound by α-bungarotoxin (**Figure 2i**), all the toxins stay bound to the nAChR throughout the simulations (**Figures S11-S12**). In an effort to determine the effects of toxin binding to the nAChR, we calculated the average root mean square fluctuations (RMSF) of the nAChR from the five simulations performed for each nAChR simulation systems. Then, we subtract the nAChR RMSF of the apo receptor from the nAChR RMSF of the systems with toxins bound to determine the changes in receptor flexibility upon toxin binding (**Figures S13-S14**). Overall, the AC nAChR became much more rigid, especially at the orthosteric pockets, the extracellular pore, the subunit interfaces, and the interface between the ligand binding domain and transmembrane helix (**Figure S13**), which were the receptor regions that were identified to bind toxins in this and previous study^46^. In addition to these regions, the α-bungarotoxin also made the transmembrane helical domains of the human α7 nAChR much more rigid, especially at the allosteric sites (**Figure S14**) identified previously^46^. Therefore, toxin binding to the nAChR appeared to rigidize all the possible binding sites in the nAChR. Furthermore, since nAChRs belong to the ligand-gated ion channel family^46^, we hypothesize that the rigidity caused by the toxins will restrict the necessary movements required by the nAChRs to open their pores for the ions to pass through, leading to toxicity.

To confirm our hypothesis that the binding to toxin modulates the gating mechanism, we performed principal component analysis (PCA) to determine the primary motions of the nAChR simulation systems in the presence and absence of the toxins (**Figure 5**). In the apo AC nAChR, the primary motions involved major outwards movements of the extracellular pore, the outward movements of the loop regions and the subunit interfaces in the nAChR orthosteric pockets, and the inward movements of the loop regions towards the subunit interfaces in the nAChR orthosteric pockets (**Figure 5a**). Similar primary motions were observed in the apo human α7 nAChR, and we also observed minor movements at the horizontal helices located at the interface of the transmembrane helical and intracellular domains (**Figure 5h**). Upon binding of the toxins to the nAChRs, the nAChR orthosteric pockets were rigidized, and no primary motion was observed at the nAChR orthosteric pockets (**Figure 5b-5g** and **5i**). However, we observed major closing motions at the extracellular pores of the soluble ligand binding domains and the horizontal helices at the interface of the transmembrane helical and intracellular domains, the two regions known to serve as the gates for the entrance of ions through the nAChRs^26^. In particular, we observed two opposite subunits moving towards another to close the extracellular pores in the AC nAChRs bound by α-conotoxin BuIA (**Figure 5b**), strychnine (**Figure 5e**), and pinnatoxin A (**Figure 5f**). The horizontal helices at the interface of the transmembrane helical and intracellular domains were observed to move strongly towards their neighbors in the presence of the α-bungarotoxin in the human α7 nAChR (**Figure 5i**). Therefore, through the closing gating mechanism, the toxins enacted their toxicity in the nAChRs by preventing the ions from diffusing through the channels^26^.

**Figure 5.**
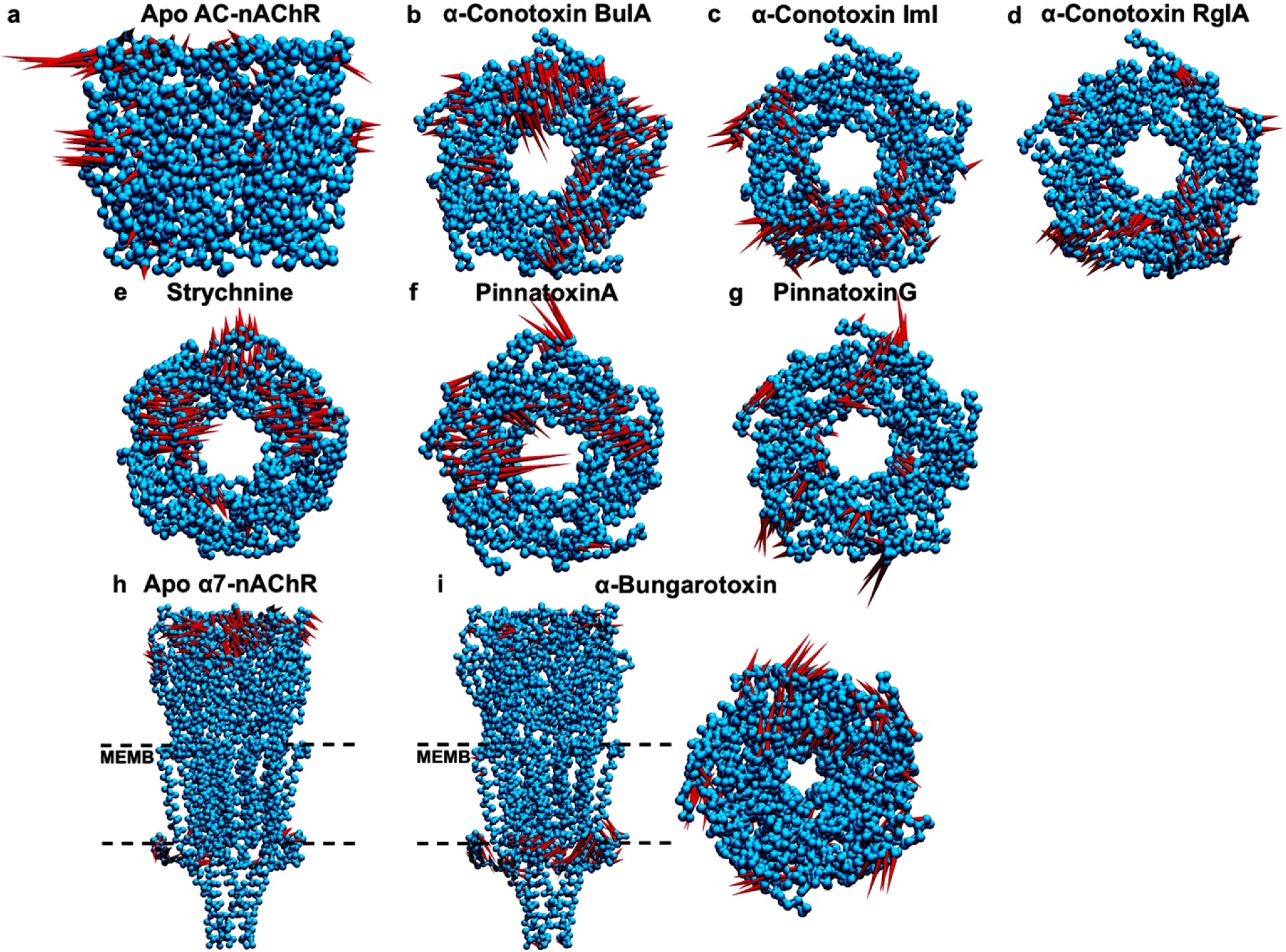
Primary motions detected from the principal component analyses of the GaMD simulations of. **(a)** apo AC nAChR, **(b)** AC nAChR bound by α-conotoxin BuIA, **(c)** AC nAChR bound by α-conotoxin ImI, **(d)** AC nAChR bound by α-conotoxin RgIA, **(e)** AC nAChR bound by strychnine, **(f)** AC nAChR bound by pinnatoxin A, **(g)** AC nAChR bound by pinnatoxin G, **(h)** apo human α7 nAChR, and **(i)** human α7 nAChR bound by α-bungarotoxin. The arrow lengths illustrate the qualitative extents of the motions. The transmembrane-bound domains in **(h)** and **(i)** were indicated by the two dashed lines (MEMB).

## Discussion

While it is known that various ligands, including toxins of different physical and structural properties, bind to the same orthosteric pockets in the nAChRs, the molecular reason behind that remains unknown. Furthermore, the binding and dissociation pathways of toxins to the nAChRs and the associated molecular descriptors have been unexplored due to the extensive computational resources required, which hinders the efforts to develop effective antidotes for the toxins. In this work, we have performed extensive all-atom dual-boost GaMD simulations on a number of nAChR systems with toxins, including α-conotoxins BuIA, ImI, and RgIA, strychnine, pinnatoxin A, pinnatoxin G, and α-bungarotoxin for a total of 60 μs of simulation time to determine the binding and dissociation pathways of toxins from the nAChRs as well as the effect of toxin binding to the receptor.

We captured two common binding and dissociation pathways that were shared across the toxins (α-conotoxin RgIA and strychnine), organisms (*Aplysia californica* and humans), and nAChR subunits (AC, human α7, and human α3β4). In the primary binding pathway of toxins to the nAChR, the toxins gradually diffused from the bulk solvent to first bind near the extracellular pore of the ligand binding domain of the nAChR, then moved downwards along the nAChR to the orthosteric pockets (**Figures 4a** and **S8-S10**). In the secondary binding pathway, the toxins diffused directly from the bulk solvents into the nAChR orthosteric pockets (**Figure S5**). The dissociation pathways of the toxins from the nAChR were the direct reverse of the observed binding pathways (**Figures 3a, S3,** and **S8b**). The secondary binding and dissociation pathways that we observed for toxins in the nAChRs were found to be highly similar to the binding pathway of the γ-aminobutyric acid (GABA) neurotransmitter to the GABA type A receptors identified from a previous study^47^. We also observed a minor binding pathway, where the toxins moved along the interface between two subunits to bind the orthosteric pocket (**Figure S9a**). Furthermore, we determined that toxin binding to the nAChRs rigidized the nAChRs, especially at the binding sites where the orthosteric and allosteric ligands could bind (**Figures S13-S14**)^46^. Based on the primary motions detected from our PCA, we concluded that the toxins enact their toxicity by restricting the gating mechanism of the extracellular pores and the horizontal helices at the interface of the transmembrane helical and intracellular domains in the nAChRs and preventing the ions from entering the nAChRs^26^.

To explain the common binding and dissociation pathways observed across different toxins and nAChRs, we calculated the electrostatic potentials ^48^ for the nAChRs and protein toxins (α-conotoxins BuIA, ImI, and RgIA and α-bungarotoxin) (**Figures 6** and **S15**). First, we observed that the orthosteric pockets of nAChR were electrostatically bipolar: the inner side of the loop that connected the two β-strands was electronegatively charged, while its opposite side to the right (located at the interface of the subunits) was mostly electropositive (**Figure 6a**). Therefore, this binding pocket was highly suitable to accommodate toxins such as α-conotoxins BuIA, ImI, and RgIA and α-bungarotoxin, with their regions of contacts to the binding pocket also being bipolar (**Figure 6b-6d**, and **S15**). The electropositively charged side of the toxins could stably interact with the inner side of the nAChR loop, while the electronegatively charged side could face the electropositive side of the nAChR orthosteric pocket (**Figure 6**). This phenomenon was also observed for the endogenous ligand of the nAChR, acetylcholine (**Figure S16**). In particular, acetylcholine also had a positively charged tail (-N(CH_3_)_3_^+^) and an electronegative end (-O-C(=O)-CH_3_). The orientation of acetylcholine was the exact same as the toxins, with the positively charged end favorably interacting with the nAChR loop, while the electronegative end interacting with the opposite site (**Figure S16**). Overall, this determinant has been proposed to be important for toxin/ligand binding in the nAChR and other neuronal receptors^47,49^.

**Figure 6.**
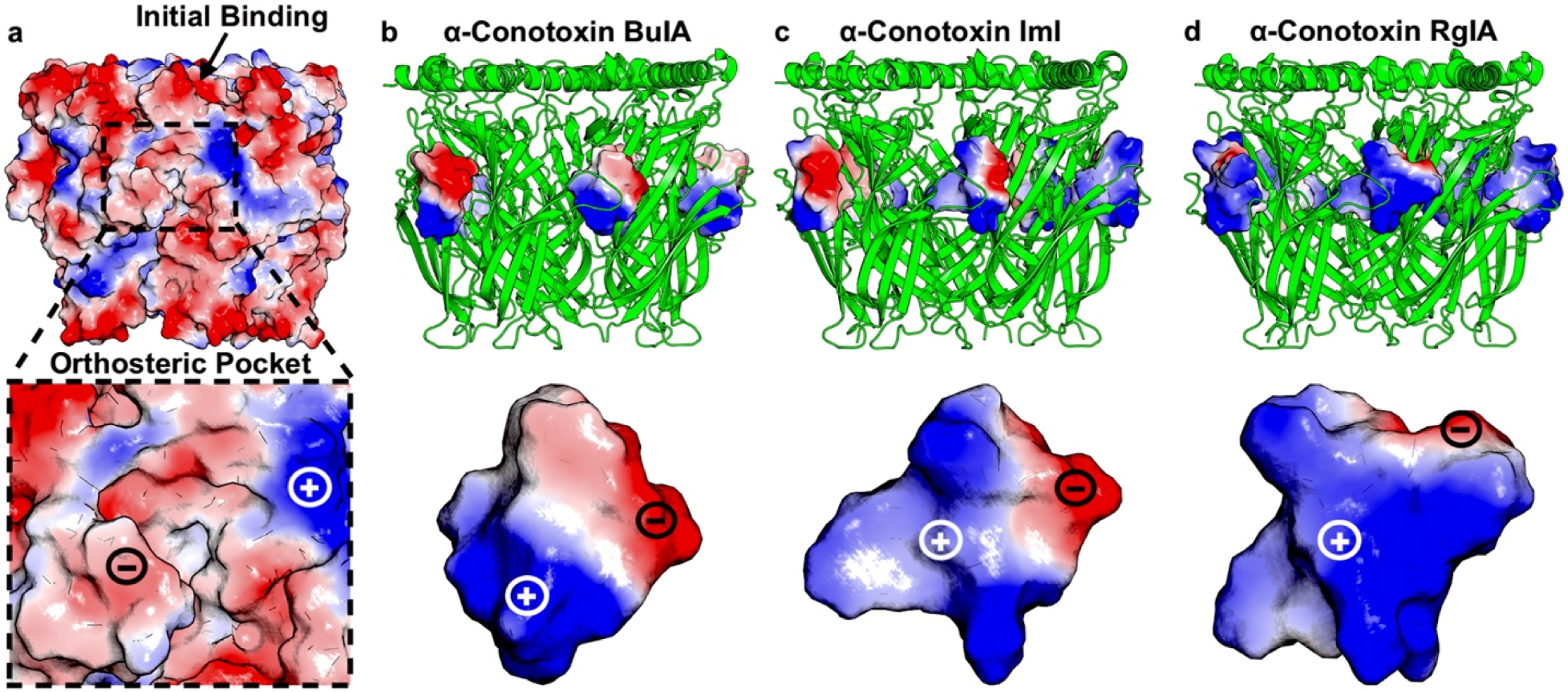
Electrostatic potential surfaces of the. Aplysia californica (AC) nicotinic acetylcholine receptor (nAChR), **(b)** α-conotoxin BuIA that binds AC nAChR, **(c)** α-conotoxin ImI that binds AC nAChR, and **(d)** α-conotoxin RgIA that binds AC nAChR. Zoom-in views of the nAChR orthosteric pocket and α-conotoxins showed high complementarity in the electrostatic potentials between the nAChR orthosteric pocket and toxins **(a-d)**. The initial binding sites of the toxins at the extracellular pore of the nAChR (indicated by an arrow in **(a)**) could only accommodate the electropositive surface of the toxins. A color scheme of red (negative) – white – blue (positive) is used to show the electrostatic potential. The electrostatic potential surfaces of the human α7 nAChR and α-bungarotoxin were shown in **Figure S15**.

Next, we examined the relationship between the binding led by the electrostatically bipolar interactions and toxicity. Since the toxins were larger and carried more electrostatic potentials on them, they could easily displace the endogenous ligand to carry out their toxicity effects. Notably, since this bipolarity could only be observed at the orthosteric pocket of the nAChR, it was the only place that could serve as the orthosteric pocket to stably accommodate the toxins and orthosteric ligands (**Figure 6a**). On the other hand, since the regions near the extracellular pores and intracellular pores were mostly electronegatively charged, they could only serve as points of contact in the binding pathways of the toxins to the nAChR orthosteric pockets, where the bipolarity of the toxins could be fully accommodated (**Figure 6a**). Furthermore, as this bipolarity extent decreased from α-conotoxin BuIA to ImI to RgIA (with the electropositive charge becoming more dominant) (**Figure 6b-6d**), the extent of closing of the gating motion caused by toxin binding also decreased in the same direction (**Figure 5b-5d**), with the α-conotoxin BuIA being most bipolar and toxic. This comparison can also be made for pinnatoxin A versus pinnatoxin G. As pinnatoxin A had an additional -COOH group compared to the -CH=CH2 group of pinnatoxin G (**Figure 1e-1f**), pinnatoxin A was more bipolar and toxic, causing much larger gate-closing movements at the extracellular pore than pinnatoxin G (**Figure 5f-5g**). Therefore, we hypothesize that the electrostatic potentials (particularly charge complementarity), in addition to other factors such as shape complementarity, plays a crucial role in the binding of toxins and exerting their toxic effects on the nAChRs.

In conclusion, we have uncovered general binding pathways of the toxins to the nAChRs and found that the electrostatic potentials played an important role in explaining the common binding pockets and binding pathways of toxins to the nAChRs. Based on our findings, we predict that drugs that are designed with proper shapes and to be highly electrostatically bipolar could serve as potential antidotes for α-conotoxins and many other toxins in general as they could bind tightly to the toxins. Furthermore, the molecular descriptors we uncovered here, including the electrostatic potentials, could serve as effective inputs to train predictive machine learning models for toxin-target binding and toxicity predictions. We intend to extend the current study, which provided important mechanistic insights into the binding of diverse toxins to the nAChRs, to include much more toxins and different subunits of the nAChRs. Given the limited availability of experimental toxin-bound nAChR structures, these models are expected to augment structure and binding prediction tools such as AlphaFold2^50^ and RosetTTAFold^51^.

## Acknowledgements

This work was supported by the Defense Threat Reduction Agency (award no. HDTRA1345035 to J.K.S.). This work was performed at the Los Alamos National Laboratory, which is operated by Triad National Security, LLC, for the National Nuclear Security Administration of the U.S. Department of Energy (contract 89233218CNA000001). The views expressed in this article are those of the authors and do not reflect the official policy or position of the U.S. Department of Defense or the U.S. Government. The authors would like to thank Daphne Stanley for her support of this work.

---

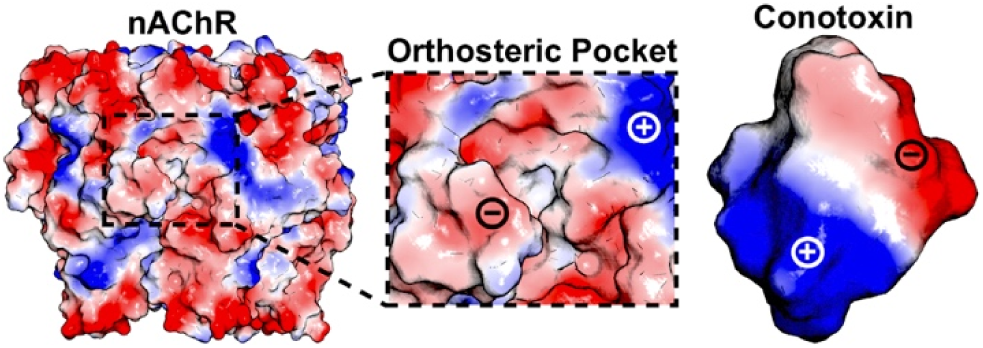
Table of Contents Graphic.

