## Supporting Information for "Diverse toxins exhibit a common binding mode to the Nicotinic Acetylcholine Receptors revealing a new molecular determinant"

**Table S1. Summary of the MD simulations of toxin binding to the nicotinic acetylcholine receptors.** AC stands for the Aplysia californica organism.

| **Simulation System** | **Initial PDB** | **Simulation Length** | **Boost Potential (kcal/mol)** |
| --- | --- | --- | --- |
| Apo AC-nAChR | 2BYN | 5 x 1µs | 12.1 ± 3.9 |
| BuIA-bound AC-nAChR | 4EZ1 | 5 x 1µs | 12.1 ± 3.9 |
| ImI-bound AC-nAChR | 2BYP | 5 x 1µs | 13.1 ± 4.0 |
| RgIA-bound AC-nAChR | 7EGR | 5 x 1µs | 12.5 ± 3.9 |
| Strychnine-bound AC-nAChR | 2XYS | 5 x 1µs | 13.5 ± 4.1 |
| PinnatoxinA-bound AC-nAChR | 4XHE | 5 x 1µs | 11.9 ± 3.8 |
| PinnatoxinG-bound AC-nAChR | 4XK9 | 5 x 1µs | 12.6 ± 3.9 |
| AC-nAChR with free RgIA | 7EGR | 5 x 1µs | 11.8 ± 3.8 |
| Apo human α7 nAChR | 7EKI | 5 x 1µs | 13.9 ± 4.6 |
| α-Bungarotoxin-bound α7 nAChR | 7KOO | 5 x 1µs | 14.4 ± 4.6 |
| α7 nAChR with free RgIA | 7EKI | 5 x 1µs | 13.5 ± 4.3 |
| α3β4 nAChR with free RgIA | 6PV7 | 5 x 1µs | 14.6 ± 4.5 |

**Figure S1. Simulation systems of α-conotoxin RgIA binding to nicotinic acetylcholine receptors (nAChRs). (a)** Aplysia californica (AC) nAChR with free RgIA α-conotoxin peptides. **(b)** Human α7 nAChR with free RgIA α-conotoxin peptides. **(c)** Human α3β4 nAChR with free RgIA α-conotoxin peptides. The nAChRs are colored green, while the toxins are colored orange. The POPC membrane lipid bilayers were shown in **(b-c)**. The PDB IDs of the structures are included in **Table S1**.

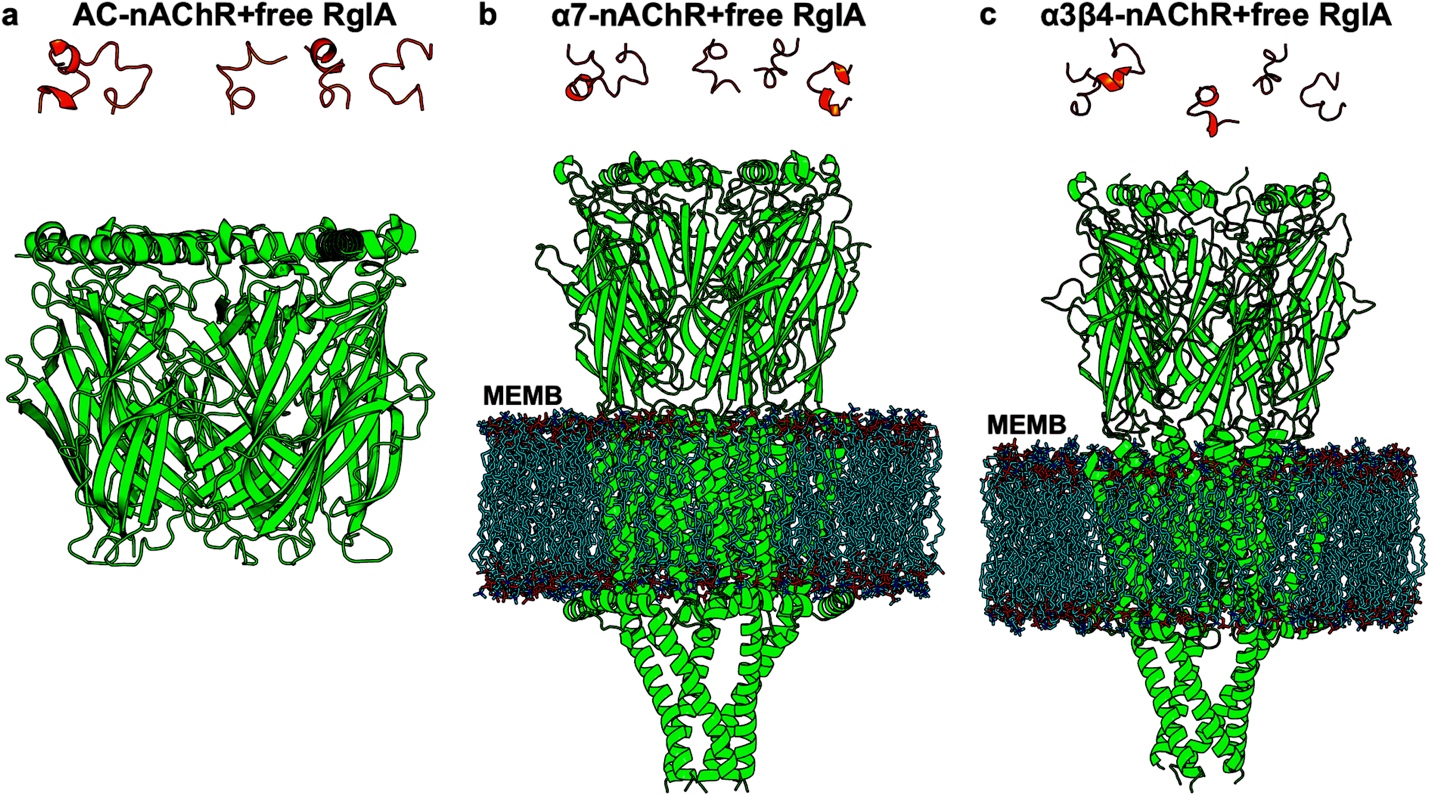

**Figure S2. Time courses of the COM distances between strychnine (excluding strychnine-2) and the Cα atoms of Aplysia californica nAChR orthosteric pockets (including residues Q186 – Y195 in the subunits) and nAChR pore diameters (Cα-atom distance between nAChR residues A:Q100 and D:Q100) calculated from the GaMD simulations of strychnine-bound AC-nAChR.**

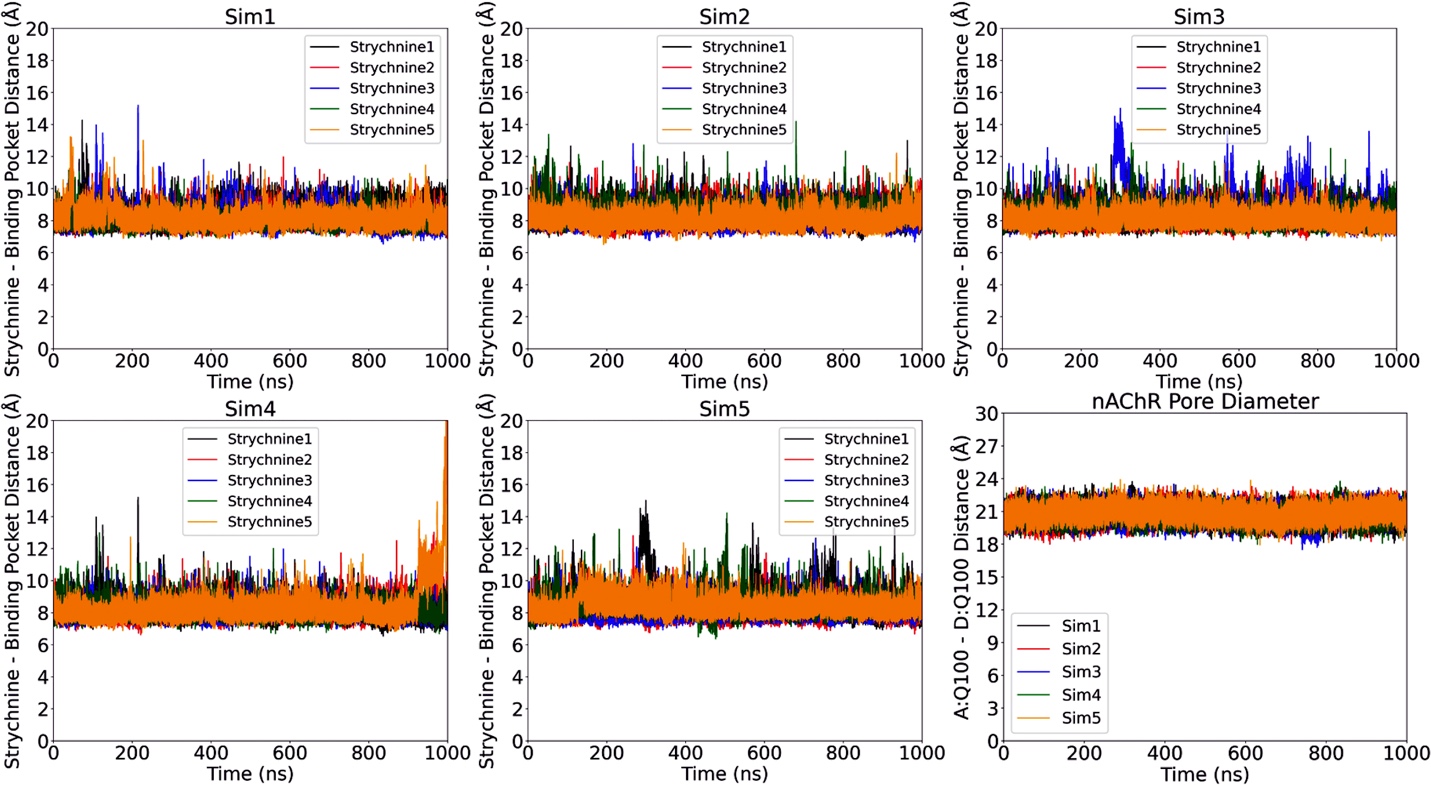

**Figure S3. Dissociation of strychnine-2 from the Aplysia californica nAChR. (a)** Time courses of the COM distances between strychnine-2 and the Cα atoms of AC nAChR orthosteric pocket (including residues E:Q186 – E:Y195) calculated from the GaMD simulations of strychnine-bound AC nAChR. **(b)** Trace of strychnine-2 dissociation from the AC nAChR observed in Sim4, where strychnine-1 dissociation was observed towards the end of the simulation.

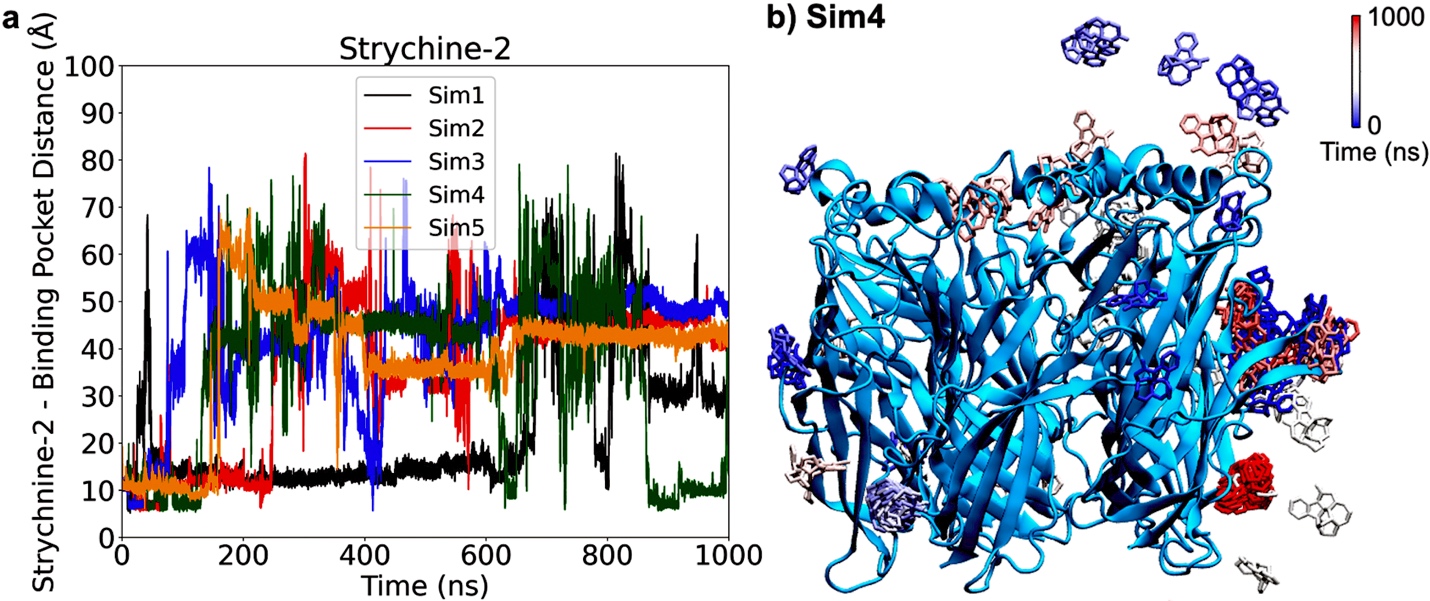

**Figure S4. Time courses of the COM distances between Cα atoms of the α-conotoxin RgIA and Aplysia californica nAChR orthosteric pockets (including residues Q186 – Y195 in the subunits) in the seven binding events and nAChR pore diameters (Cα-atom distance between nAChR residues A:Q100 and D:Q100) calculated from the GaMD simulations of AC-nAChR with free RgIA.**

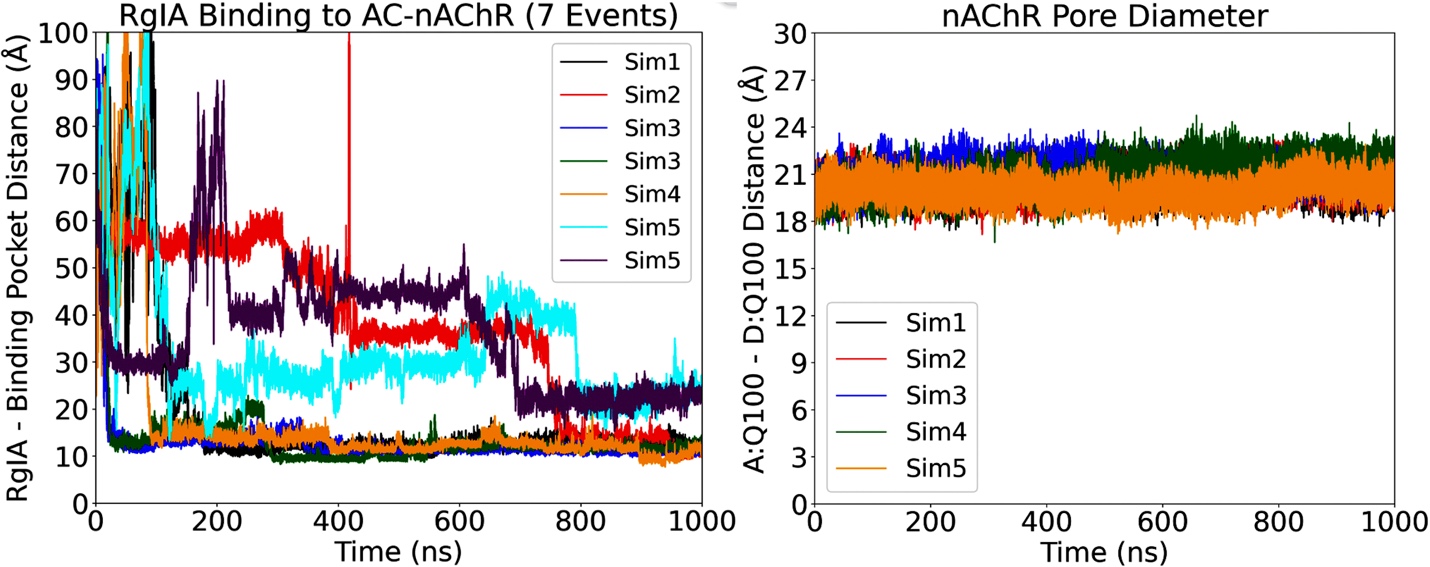

**Figure S5. Trace of α-conotoxin RgIA binding to the Aplysia californica nAChR observed in Sim1.** The binding pathway is different from what is shown in **Figure 3a**. A color scale of blue (0 ns) – white – red (1000 ns) is used to show the location of α-conotoxin RgIA during Sim1.

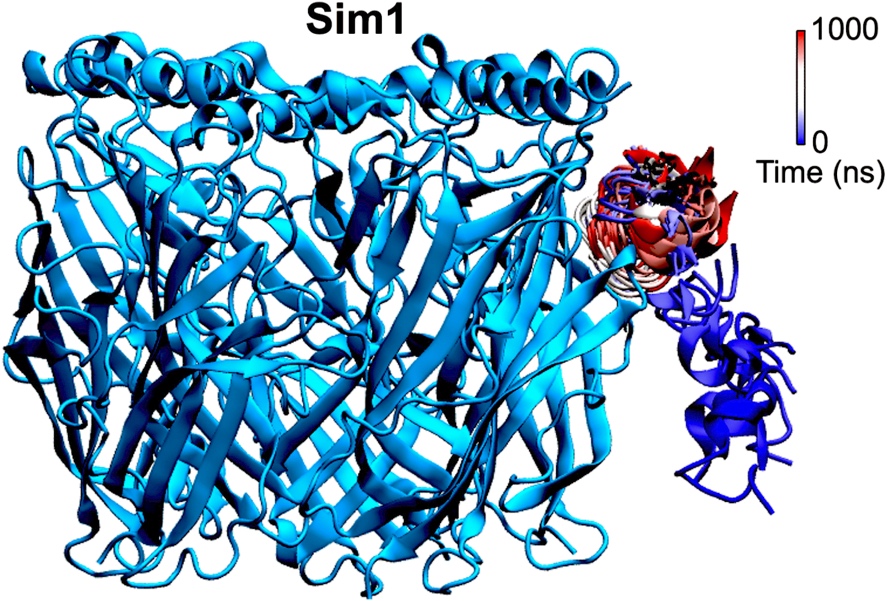

**Figure S6. Time courses of the COM distances between Cα atoms of the α-conotoxin RgIA and human α7-nAChR orthosteric pockets (including residues R185 – Y194 in the subunits) in the eight binding events and nAChR pore diameters (Cα-atom distance between nAChR residues A:A101 and D:A101) calculated from the GaMD simulations of the human α7-nAChR with free RgIA.**

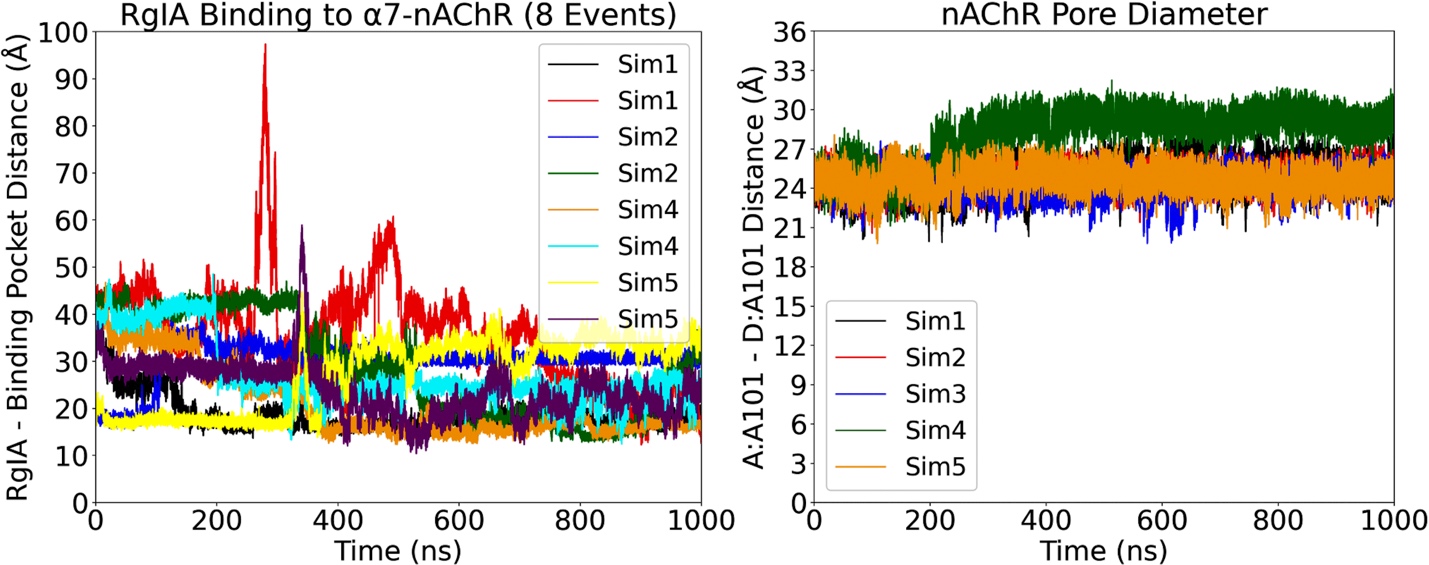

**Figure S7. Time courses of the COM distances between Cα atoms of the α-conotoxin RgIA and human α3β4-nAChR orthosteric pockets (including residues A:I188 – A:Y197, B:T190 – B:D200, C:T190 – C:D200, D:I188 – D:Y197, and E:T190 – E:D200) in the four binding events and nAChR pore diameters (Cα-atom distance between nAChR residues A:D103 and C:T103) calculated from the GaMD simulations of the human α3β4-nAChR with free RgIA.**

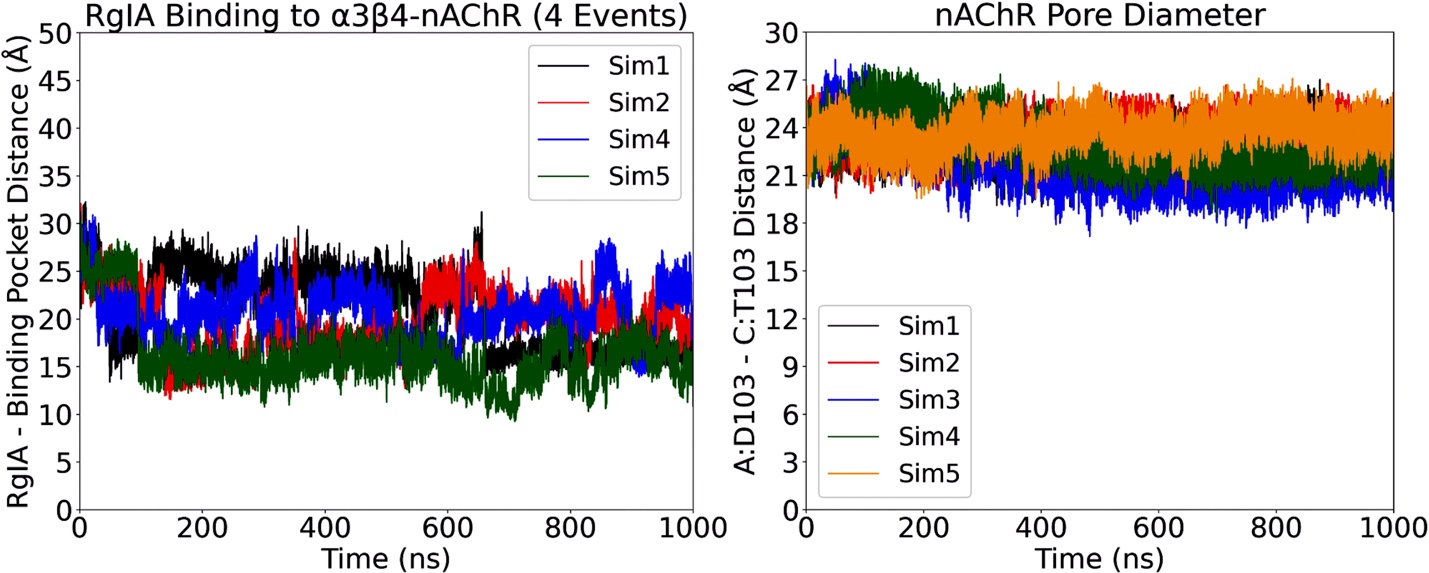

**Figure S8. Traces of α-conotoxin RgIA binding and dissociation in the human α7-nAChR observed in GaMD equilibration (a) as well as Sim2 and Sim5 (b).** A color scale of blue (0 ns) – white – red (50/1000 ns) is used to show the location of α-conotoxin RgIA during the 50 ns GaMD equilibration **(a)** as well as 1000ns Sim2 and Sim5 **(b)**, respectively.

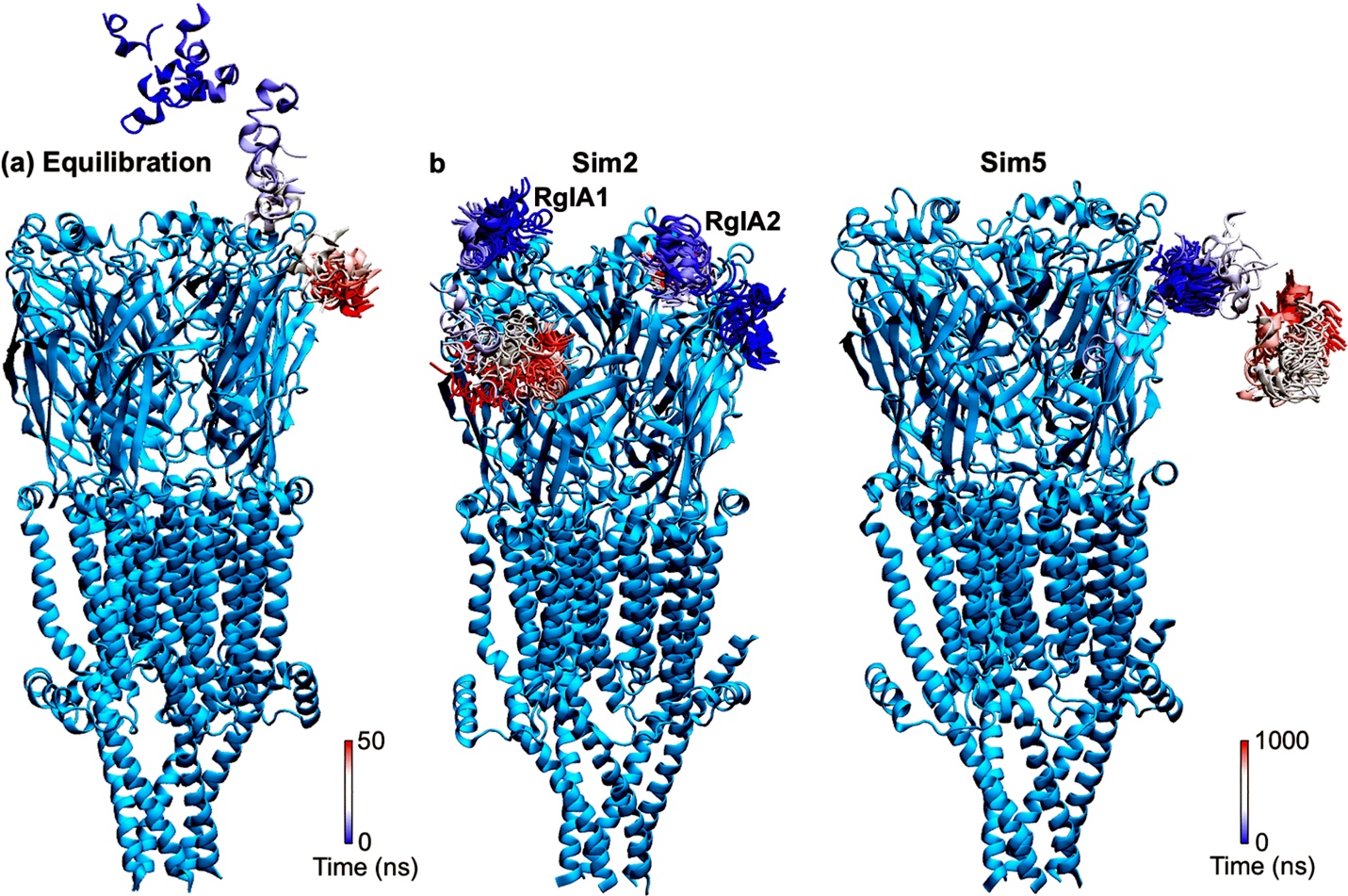

**Figure S9. Binding of α-conotoxin RgIA to the human α7 nicotinic acetylcholine receptor. (a)** Representative trace of α-conotoxin RgIA binding to the α7 nAChR observed in Sim1. A color scale of blue (0 ns) – white – red (1000 ns) is used to show the location of α-conotoxin RgIA during Sim1. **(b)** 2D potential mean force (PMF) free energy profile of the α-conotoxin RgIA binding to the α7 nAChR calculated from the COM distance between Cα atoms of RgIA and the orthosteric binding pocket of α7 nAChR (consisting of residues R185 – Y194 in the subunits) and the nAChR pore diameter (Cα-distance between residues A:A101 and D:A101). Selected low-energy conformational states are labeled “S1”–“S4”. **(c)** The “S1” state, where the distance between RgIA and nAChR orthosteric pocket is ~14.1 Å and the nAChR pore diameter is ~23.7 Å. **(d)** The “S2” state, where the distance between RgIA and nAChR orthosteric pocket is ~18.3 Å and the nAChR pore diameter is ~25.0 Å. **(e)** The “S3” state, where the distance between RgIA and nAChR orthosteric pocket is ~24.7 Å and the nAChR pore diameter is ~23.9 Å. **(f)** The “S4” state, where the distance between RgIA and nAChR orthosteric pocket is ~39.4 Å and the nAChR pore diameter is ~24.0 Å. The α-conotoxin RgIA of interest is colored and circled in red, while the other α-conotoxin RgIA molecules are colored yellow.

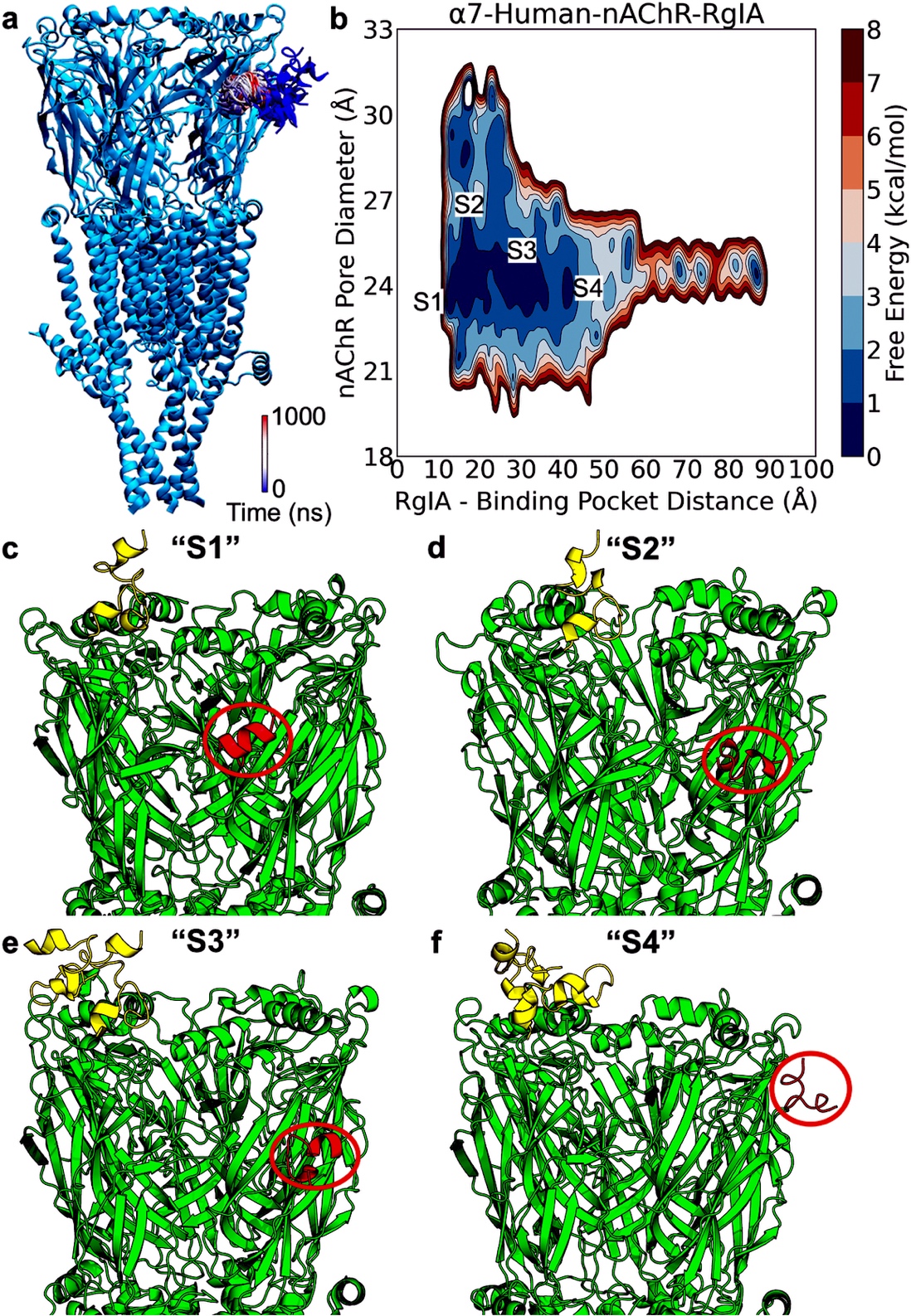

**Figure S10. Binding of α-conotoxin RgIA to the human α3β4 nicotinic acetylcholine receptor. (a)** Representative trace of α-conotoxin RgIA binding to the α3β4 nAChR observed in Sim5. A color scale of blue (0 ns) – white – red (1000 ns) is used to show the location of α-conotoxin RgIA during Sim1. **(b)** 2D potential mean force (PMF) free energy profile of the α-conotoxin RgIA binding to the α3β4 nAChR calculated from the COM distance between Cα atoms of RgIA and the orthosteric binding pocket of α3β4 nAChR (consisting of residues A:I188 – A:Y197, B:T190 – B:D200, C:T190 – C:D200, D:I188 – D:Y197, and E:T190 – E:D200) and the nAChR pore diameter (Cα-distance between residues A:D103 and C:T103). Selected low-energy conformational states are labeled “S1”–“S3”. **(c)** The “S1” state, where the distance between RgIA and nAChR orthosteric pocket is ~9.3 Å and the nAChR pore diameter is ~23.5 Å. **(d)** The “S2” state, where the distance between RgIA and nAChR orthosteric pocket is ~15.0 Å and the nAChR pore diameter is ~22.4 Å. **(e)** The “S3” state, where the distance between RgIA and nAChR orthosteric pocket is ~20.1 Å and the nAChR pore diameter is ~21.4 Å. The α-conotoxin RgIA of interest is colored and circled in red, while the other α-conotoxin RgIA molecules are colored yellow.

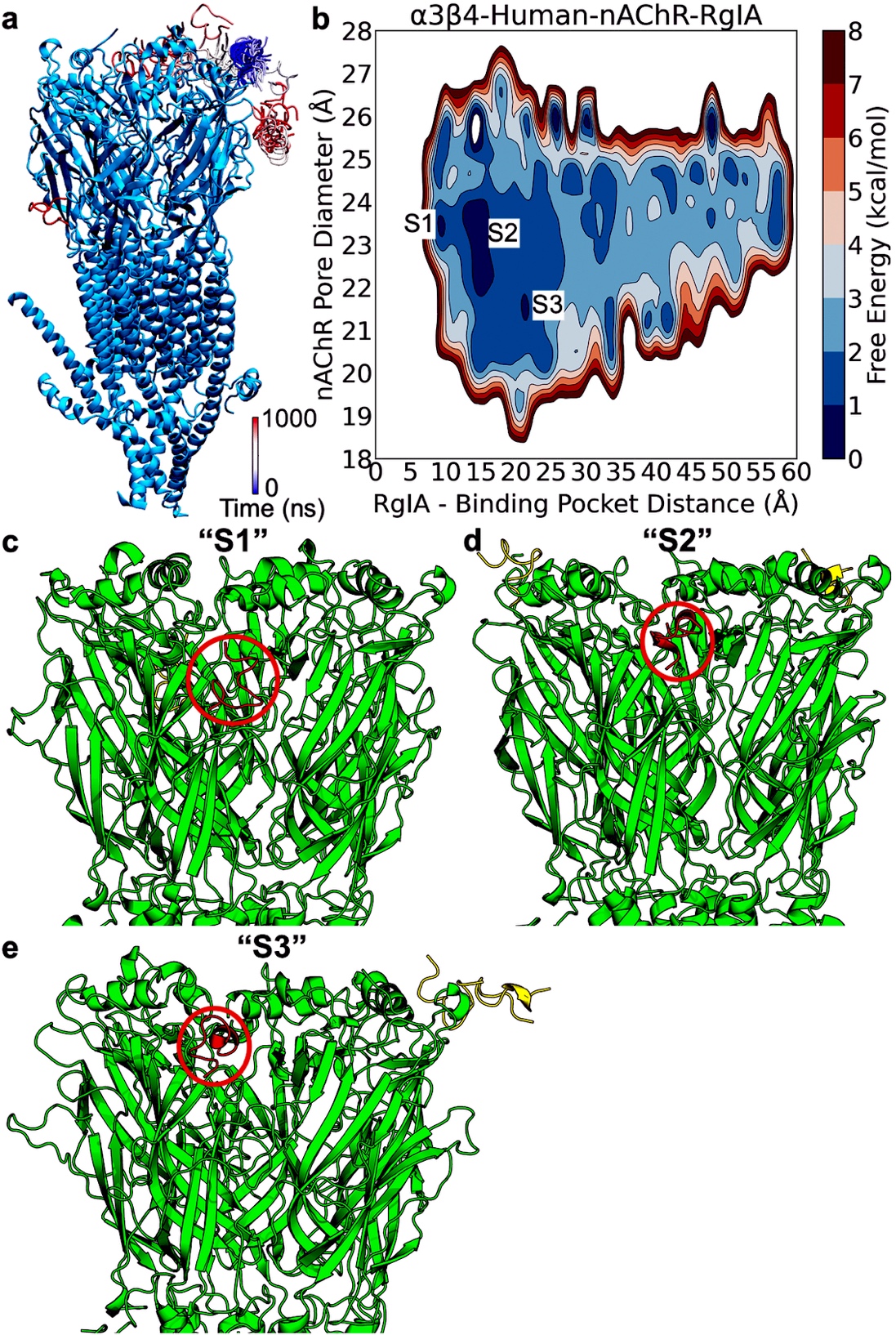

**Figure S11. Representative time courses of the COM distances between (Cα atoms of) toxins and the AC-nAChR orthosteric pockets (including residues Q186 – Y195 in the subunits) and nAChR pore diameters (Cα-atom distance between nAChR residues A:Q100 and D:Q100) calculated from the GaMD simulations of the apo and toxin-bound Aplysia californica nAChR (except strychnine).**

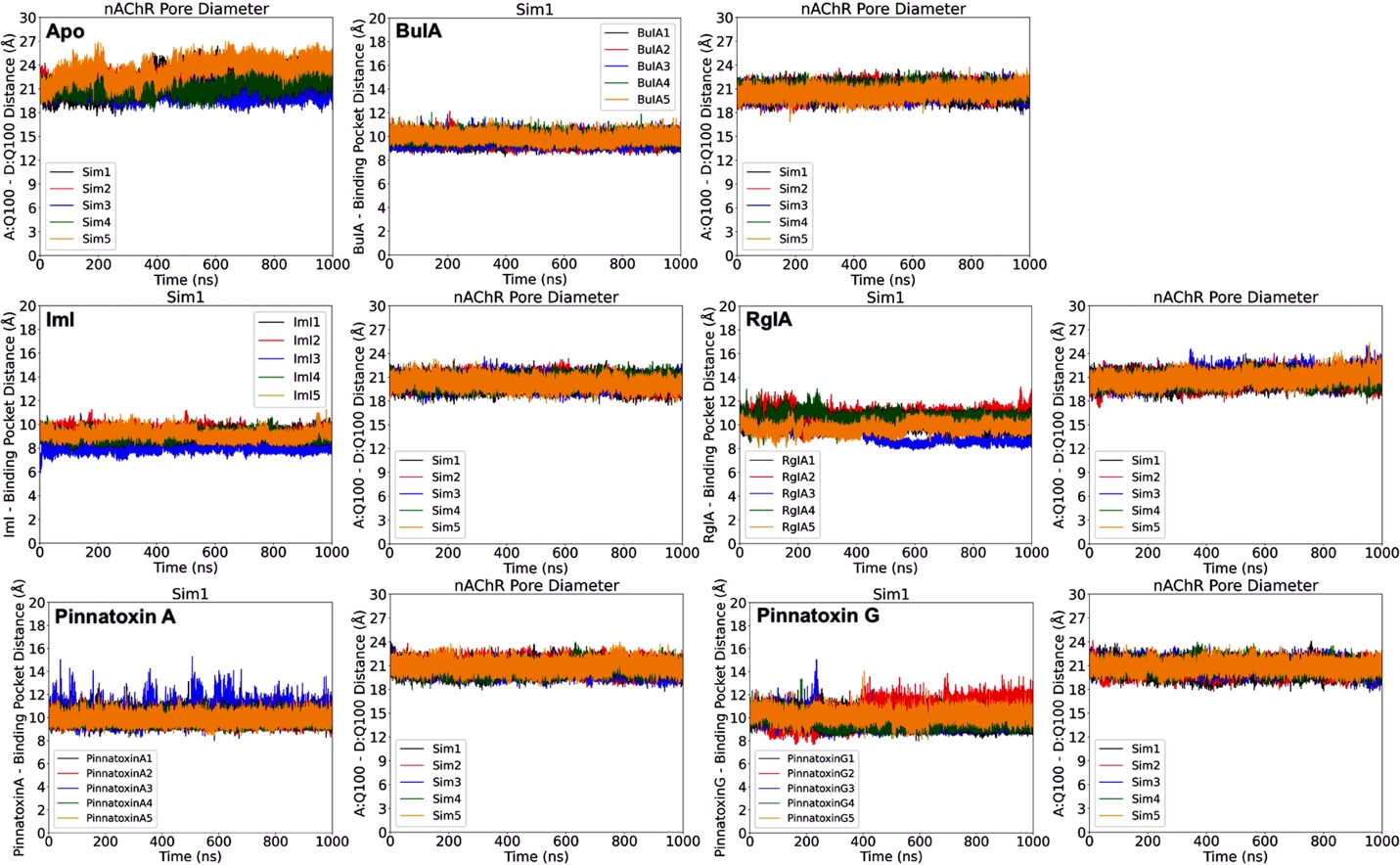

**Figure S12. Representative time courses of the COM distances between Cα atoms of the α-bungarotoxin and human α7-nAChR orthosteric pockets (including residues R185 – Y194 in the subnits) and nAChR pore diameters (Cα-atom distance between nAChR residues A:A101 and D:A101) calculated from the GaMD simulations of the apo and α-bungarotoxin bound human α7-nAChR.**

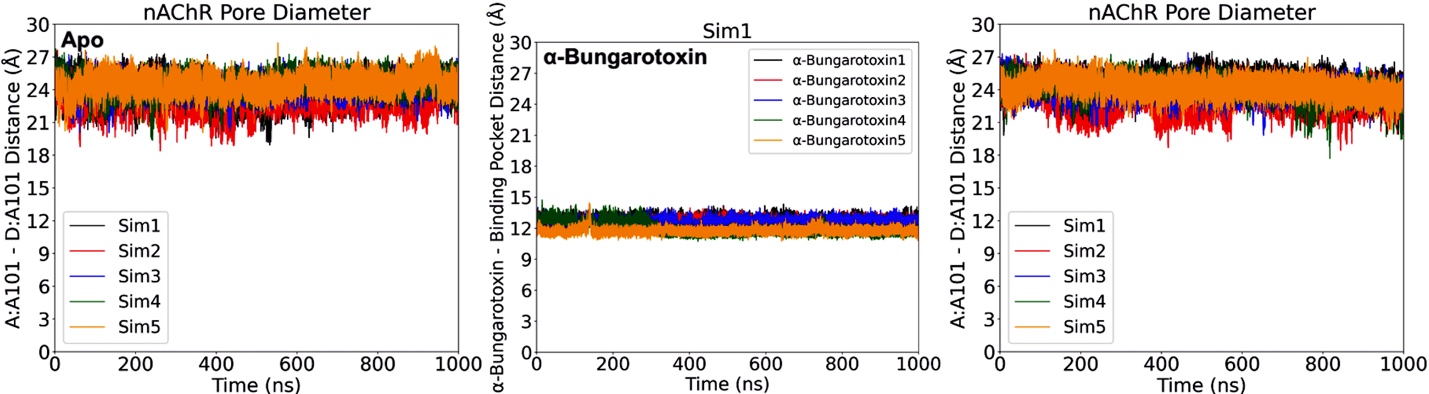

**Figure S13. Changes in the nAChR flexibility upon toxin binding calculated from the GaMD simulations of the apo and toxin bound Aplysia californica nAChR.** A color scale of blue (-0.5 Å) – white – red (0.5) is used to show the changes in receptor flexibility upon toxin binding using the root mean square fluctuation (RMSF) as metrics.

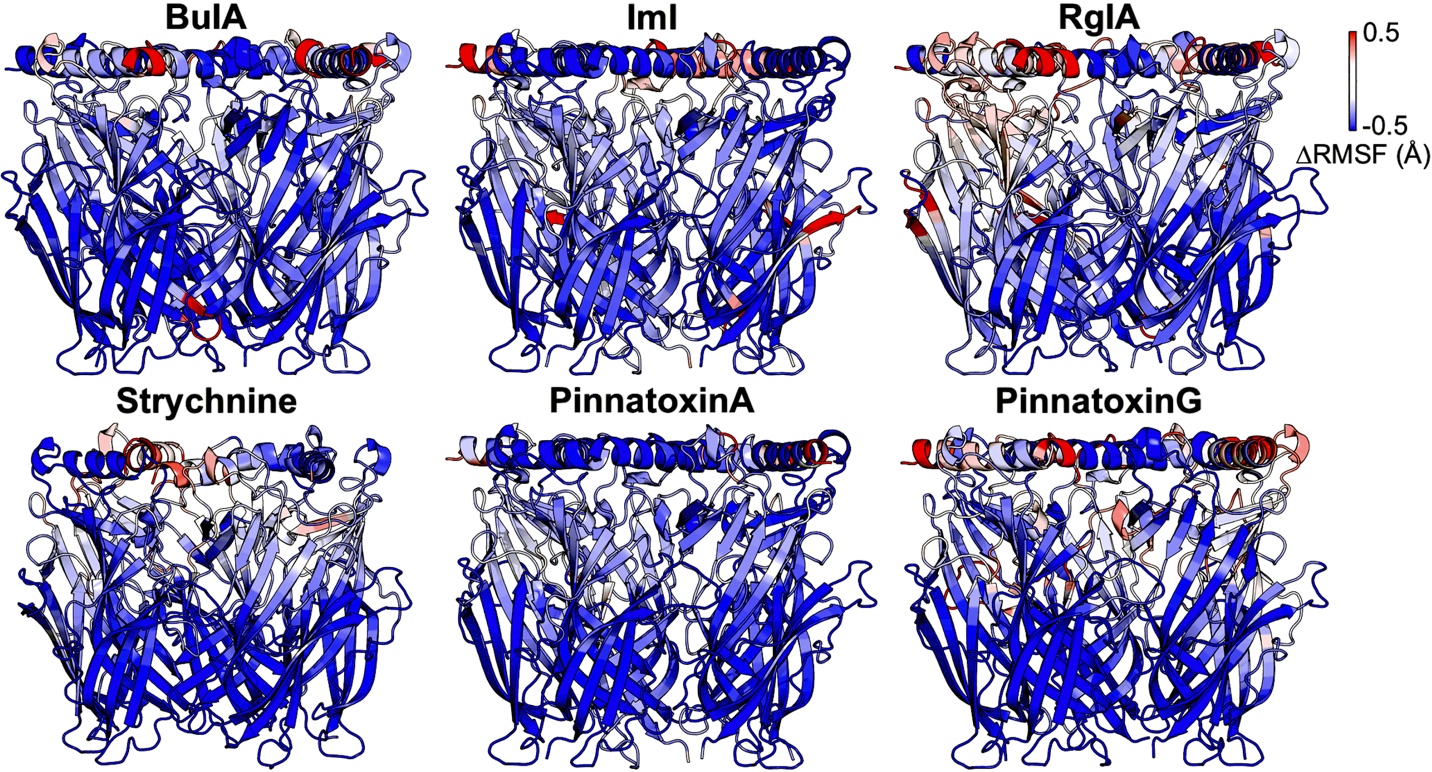

**Figure S14. Changes in the nAChR flexibility upon toxin binding calculated from the GaMD simulations of the apo and α-bungarotoxin bound human α7 nAChR.** A color scale of blue (-0.5 Å) – white – red (0.5) is used to show the changes in receptor flexibility upon toxin binding using the root mean square fluctuation (RMSF) as metrics.

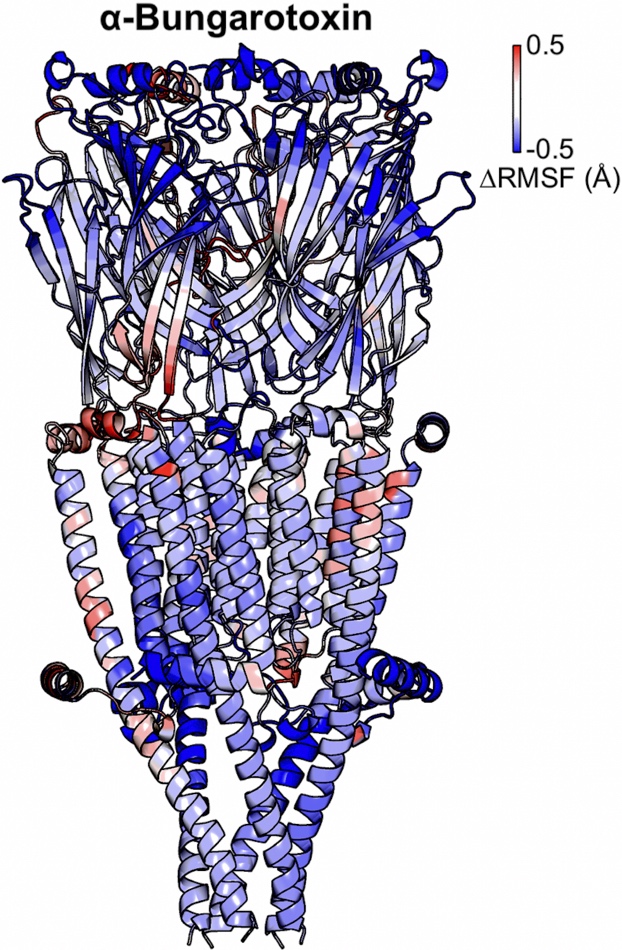

**Figure S15. Complementarity in electrostatic potential observed in the binding of α-bungarotoxin to the human α7 nicotinic acetylcholine receptor.** A color scheme of red (negative) – white – blue (positive) is used to show the electrostatic potential.

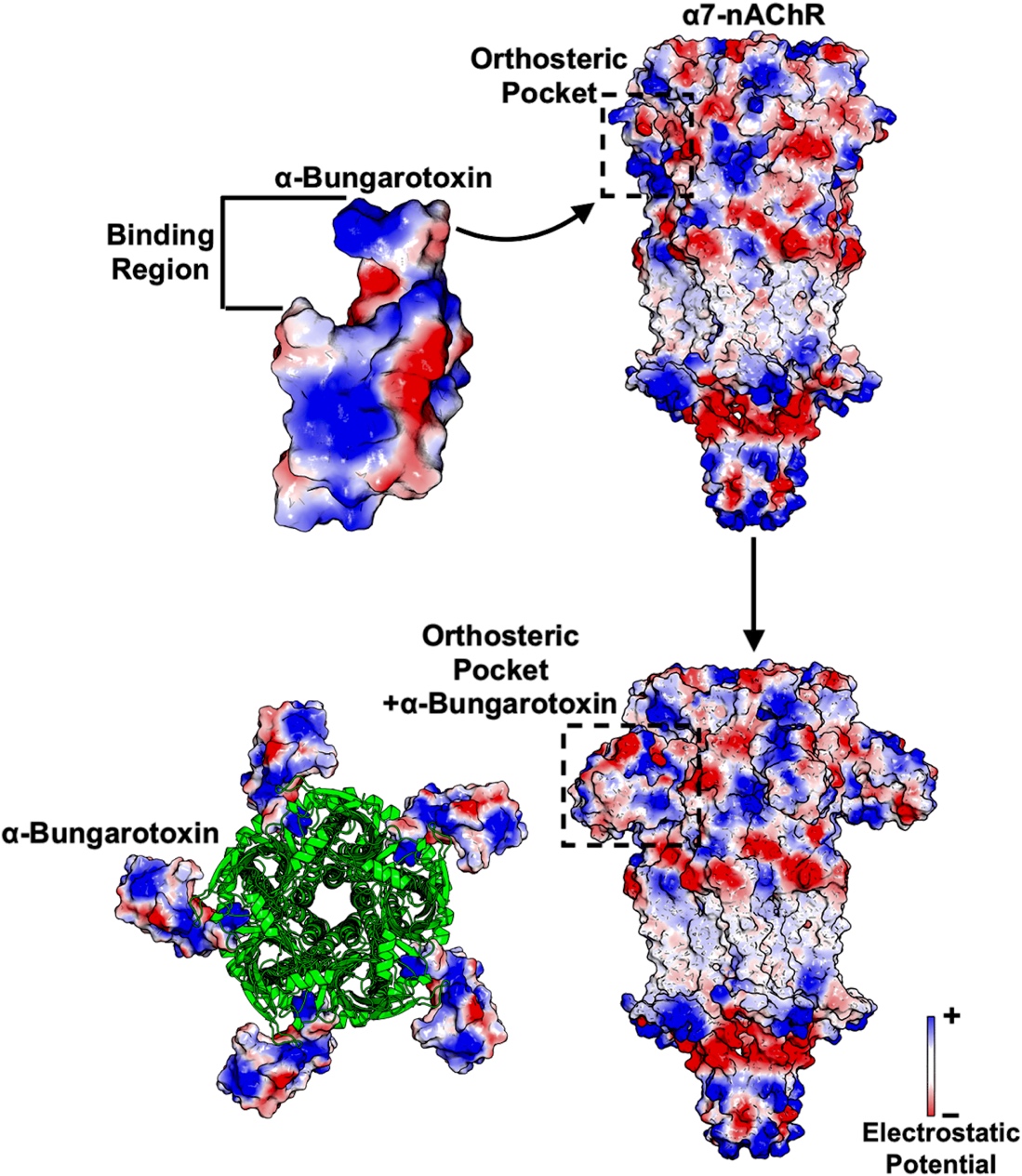

**Figure S16. Binding pose of the endogenous ligand acetylcholine in the nicotinic acetylcholine receptor.**

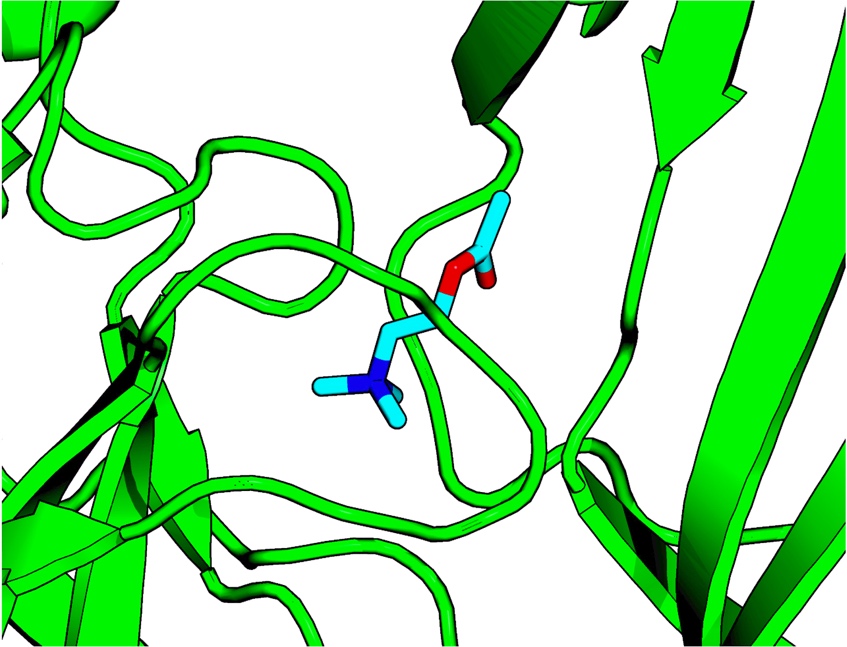
